# *MetAR*: A semi-automated meta-analysis of skeletal muscle androgen receptors association with age

**DOI:** 10.64898/2026.04.21.719741

**Authors:** Ross M. Williams, Viktor Engman, Megan Soria, Glenn Wadley, Danielle Hiam, Séverine Lamon

## Abstract

Ageing is associated with reduced skeletal muscle mass and function. Testosterone, the main androgenic hormone, exerts its anabolic effects in muscle through androgen receptor (AR) mediated pathways, with evidence suggesting that AR availability may represent a rate-limiting factor in androgen signalling. Rodent studies report age-related reductions in AR expression, however findings in human muscle are inconclusive. This study aimed to characterise age-related changes in AR expression and its upstream and downstream regulators in human skeletal muscle. To this end, we developed a novel R-based pipeline, *MetAR,* for reproducible meta-analyses of publicly available RNA sequencing (RNA-seq) datasets, to investigate associations between target gene expression and variables of interest. Eligible datasets included skeletal muscle samples from healthy adult males aged ≥18 years, with an age range of ≥ 10 years and sample size ≥ 6. Raw counts data were downloaded, appraised and trimmed mean of M-values (TMM) normalised. Dataset-level associations between age and target gene expression were modelled and random-effects meta-analyses performed. Sixteen RNA-seq datasets (n = 364; age 18-92 years) were included. AR expression was negatively associated with age (β = -0.006 log_2_ TMM-Counts Per Million (CPM) per year, *p* < 0.001, 4.4% decline per decade, I-squared=53.8%, 95% prediction interval [-0.0143 to +0.0014]). Age was associated with a reduction in expression of regulators of AR stability, transcriptional activity and nuclear transport, as well as downstream regulators relevant to AR signalling. These findings suggest multi-faceted age-associated remodelling of AR expression, signalling and nuclear transport, potentially explaining a component of age-associated muscle loss.

## Introduction

Skeletal muscle is an essential component of healthy living, enabling movement, locomotion, sustaining correct posture and promoting circulation (1-4). Skeletal muscle is equally central to maintaining energy homeostasis throughout the body, thereby preventing the development of insulin resistance, obesity and diabetes (5, 6). Humans naturally experience significant decrements in skeletal muscle mass and strength as they age, losing 3-8% of muscle mass per decade from the age of 40, with the rate of decline accelerating from the age of 60 (7, 8). Approximately 10% of older adults are diagnosed with sarcopenia, a more severe age- associated decline in skeletal muscle characterised by low muscle mass, strength, physical performance and intrinsic capacity (9-11). Sarcopenia induces significant decrements in quality of life, promoting cognitive decline, social isolation and a loss of independence (12-16), and imposes a substantial burden on healthcare systems (17).

In humans, testosterone promotes skeletal muscle anabolism primarily through androgen receptor (AR)-dependent mechanisms that enhance muscle protein accretion and myogenic capacity. Prior to androgen binding, the AR is maintained in a ligand-responsive state by multiprotein chaperone complex that includes HSP40, HSP70, HSP90, p23 and co-chaperones such as FKBP51 and FKBP52. The binding of testosterone or dihydrotestosterone induces conformational changes in the AR-chaperone complex, exposing the ARs nuclear localisation signal (NLS) to nuclear import proteins and promoting translocation to the cell nucleus. Within the nucleus, the AR complex binds to androgen-response elements (AREs) and co-regulators to regulate transcription (18-20).

Testosterone administration increases skeletal muscle protein synthesis and net protein balance by enhancing intracellular amino acid reutilisation (21, 22), increasing translational efficiency and capacity, including molecular programmes associated with ribosomal biogenesis and protein turnover (23), and increasing satellite-cell number, satellite-cell proliferation and myonuclear accretion in parallel with muscle-fibre hypertrophy (24, 25). Pre-clinical studies indicate that testosterone acts in skeletal muscle through both classical genomic AR signalling and rapid non-genomic mechanisms involving kinase and Ca²□signalling (26). Although activation of PI3K–Akt–mTORC1 signalling provides a plausible mechanism linking androgen signalling to increased translation, as demonstrated in rodent skeletal muscle *in vitro* models (27), direct evidence that sustained activation of this pathway is the principal driver of testosterone-induced hypertrophy in human skeletal muscle remains limited. Instead, mechanistic cell studies indicate that AR signalling interacts with β-catenin/TCF4 to promote myogenic differentiation and increases follistatin expression, thereby antagonising activin/myostatin/TGF-β–Smad signalling and removing inhibitory constraints on muscle growth (28, 29). Together, these pathways promote muscle protein synthesis, supress protein degradation and stimulate satellite cell proliferation (24, 25, 30-32).

Whilst there is some evidence for meaningful associations between endogenous testosterone and sarcopenia, inferences are limited by methodological heterogeneity and control for covariates such as testosterone replacement therapy (33-36). Other aspects of the androgen signalling cascade may however be more closely linked with the development of age-related muscle wasting and sarcopenia. Recent research has reported no correlation between serum testosterone concentrations and the level of hypertrophic response to resistance training (37). Instead, the rate-limiting factor in androgen-driven muscle growth may be saturation of the AR (37). Similar saturation models have been proposed in ARs in rat and human prostate tissue, which can become saturated below the normal physiological levels of testosterone (38). These findings suggest that physiological serum testosterone concentrations may exceed AR binding capacity. As such, reduced androgen binding due to limited receptor availability may play a role in age-related muscle wasting. In line with this hypothesis, prior research in rat prostate and penile tissue have shown age-associated decreases of up to 50% in AR mRNA and protein expression (39, 40). However, research in human muscle is limited, reporting discordant associations in small cross-sectional cohorts of males that were not specifically powered for examining age-related changes in AR expression (41-44). Critically, no study has systematically examined the effects of ageing on AR expression and its associated transcriptional machinery. Since the introduction of high-throughput -omics technologies in the late twentieth century, a wealth of data regarding the human transcriptome has been generated, with many datasets being made publicly available through repositories such as NCBI Gene Expression Omnibus (GEO). This study applies a novel meta-analytic approach to publicly available RNA-seq datasets, with the aim of characterising age-related changes in expression of the AR, its regulators and downstream regulators relevant to AR signalling in human skeletal muscle across the male lifespan.

## Methodology

We developed an original R pipeline, *MetAR*, for the purpose of this study. *MetAR* facilitates accessible, efficient and reproducible meta-analysis of publicly available RNA-seq datasets. *MetAR* downloads sample characteristics data and raw counts data from NCBI GEO, performs dataset-level pre-processing and statistical modelling, and conducts a random effects meta-analysis of the effect of a variable of interest on target gene expression. This study used *MetAR* (version 1.0), available at: https://github.com/ROSWILLIAM/MetAR. Prior to commencing dataset selection, the review protocol was registered with the International Prospective Register of Systematic Reviews (PROSPERO; ID: CRD420251030652).

### Dataset eligibility

The Population, Intervention, Comparison, Outcomes and Study design (PICOS) framework was used to define the dataset eligibility criteria. Eligible datasets consisted of skeletal muscle tissue from healthy biological male donors aged ≥ 18 years, with an age span of ≥ 10 between youngest and oldest donors to ensure sufficient age variation for modelling continuous age effects. Analyses were restricted to males as gene-expression in skeletal muscle is highly sex-specific, with 3000 genes being differentially expressed between sexes (45, 46). Where datasets included participants diagnosed with conditions or undergoing treatments potentially affecting skeletal muscle, hormone concentrations or AR expression (e.g. cancer, kidney disease and testosterone replacement therapy), only healthy control samples were included. This approach ensured that observed associations were not confounded by disease or treatment related changes in gene expression. Age was treated as a continuous variable and as such, comparisons were made across the age range within each dataset. Eligible outcomes were raw gene expression counts matrices from skeletal muscle genome-wide bulk RNA-sequencing, with ≥ 6 eligible participants to minimise false discovery rate (47).

### Dataset search strategy

Publicly available datasets were identified through systematic searching of NCBI GEO, using the search term “muscle”. The following search filters were applied: species = “Homo sapiens”; Supplementary files = “.txt”, “.csv”; Method = “Expression profiling by high throughput sequencing”. Final searches were conducted on 14/10/2025.

### Dataset screening

NCBI GEO dataset records were screened in duplicate by authors RMW and VE. Sample level metadata (pData) were retrieved using the r package *GeoQuery* (version 2.70.0) (48) and screened (Figure 1). Where critical data (participant age and sex) was not available, dataset custodians were contacted and offered two weeks to provide the requested data or were omitted from the final analysis.

**Figure 1.**
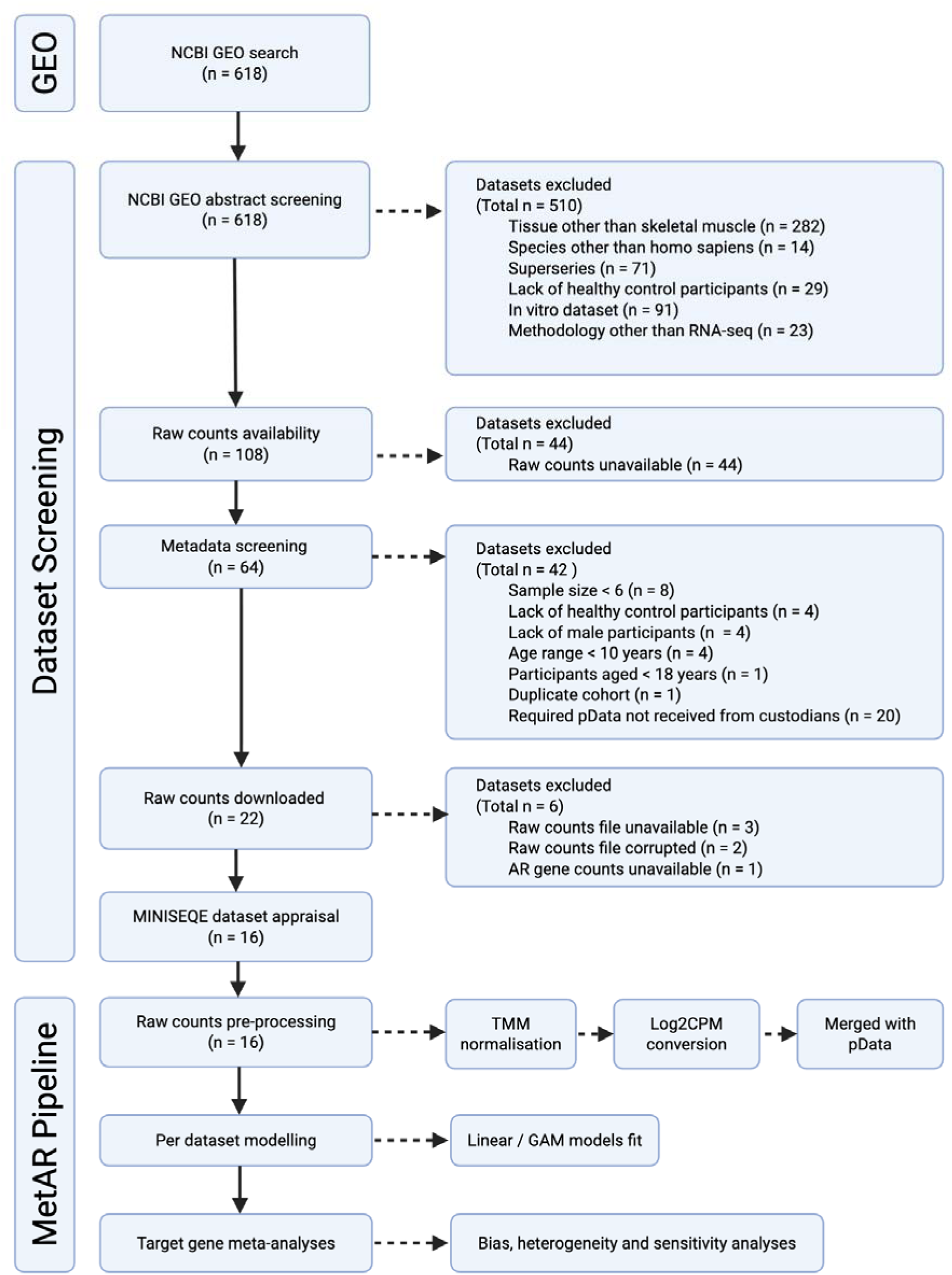
PRISMA / MetAR dataset selection and meta-analysis flowchart. *MINSEQE: Minimum Information About a Next-generation Sequencing Experiment; TMM: Trimmed mean of M-values; CPM: Counts per million; GAM: Generalised additive model. Created with BioRender.*

### Dataset quality appraisal

Eligible datasets were appraised in duplicate by authors RMW and VE in accordance with the Minimum Information About a Next-generation Sequencing Experiment (MNISEQE) guidelines (49, 50). Criteria encompassed raw data availability, sample annotation, experimental design and the documentation of essential laboratory and processing protocols. Datasets were appraised using a critical inclusion criterion and a quality index.

### Dataset pre-processing

Raw counts matrices were downloaded from NCBI GEO using *GeoQuery* and were filtered prior to normalisation using a minimal expression threshold of > 1 counts per million in at least two samples. Raw counts were normalised using the trimmed mean of M-values normalisation (TMM) to account for differences in library size and sequencing depth using the R package *edgeR* (version 4.0.16) (51, 52). Normalised counts were converted to log_2_ TMM counts per million (CPM). Per dataset, log_2_ TMM-CPM expression values were extracted from the count matrices for each target gene and merged with the corresponding processed pData.

### Meta-analysis

For each dataset, both linear and generalised additive models (GAM) were fitted to estimate the relationship between target gene expression (TMM normalised log_2_ CPM counts) and age. For linear models, the slope (β) and standard error (SE) were extracted. As GAMs capture non-linear relationships between age and gene expression, they do not provide a single slope estimate comparable to linear models therefore, the weighted mean derivative was used to summarise the average rate of change across the observed age range as a single effect size suitable for meta-analysis. For GAM models, predicted expression values were evaluated across 200 equally spaced points spanning the observed age range. The derivative of the predicted gene expression with respect to age was calculated at each point and weighted by the estimated density of observed ages, reducing bias from uneven age distributions. The weighted mean derivative was used as the GAM effect size, representing the average rate of change in gene expression per year across the observed age distribution. Standard errors were estimated using non-parametric bootstrapping. For each dataset, the data were resampled and the GAM refitted. The weighted mean derivative was recalculated for each bootstrapped sample. This was iterated 1000 times, and the standard deviation of the bootstrap effect sizes was used as the standard error for the GAM model. Model fit was assessed using estimated degrees of freedom and Akaike Information Criterion (AIC). GAM models were utilised where estimated degrees of freedom were > 1.7 and the AIC delta was ≥ 5, representing a more conservative threshold to ensure clear improvement in the model fit and stronger evidence of non-linearity.

Random-effects meta-analyses were conducted using the R package *metafor* (version 4.8.0) (53) to produce pooled estimates of the effect of age on target gene expression, based on regression coefficients (β) from linear models and weighted mean derivatives from GAM models. Models were fitted with restricted maximum likelihood (REML), which accounts for within study sampling error and between study heterogeneity. Only genes present in a minimum of 5 datasets were eligible for meta-analyses, to ensure robustness of variance estimates and reduce the influence of single studies, consistent with previously published transcriptomic meta-analyses (54, 55). Publication bias was assessed using visual inspection of funnel plots and statistically tested using Egger’s regression test and Begg’s rank test. Heterogeneity across studies was assessed using the I² statistic and Cochran’s Q test, with significance set at *p* < 0.05, and visual inspection of Baujat plots. 95% prediction intervals (PIs) were calculated to predict the range of true effects in future datasets. Sensitivity analyses were conducted using leave-one-out analysis to assess the influence of individual studies on the meta-analysis effect estimates. Significance for association between individual genes and age was set at a false discovery rate (*FDR)* < 0.05.

The results of each individual gene-level meta-analysis were tabulated (Table 1) and summarized in categories based on gene function (Table 2). Pooled effect sizes for each meta-analysis were visualized using forest plots (Figures 3-5). Results of the leave-one-out sensitivity analyses are presented in Supplementary Table 3, with influential datasets summarized in Supplementary Table 4. Due to the large number of genes analysed and figures plotted, individual Baujat plots, funnel plots and forest plots, and underlying linear and GAM model scatterplots are available at https://doi.org/10.26187/deakin.31920696.

**Table 1.**
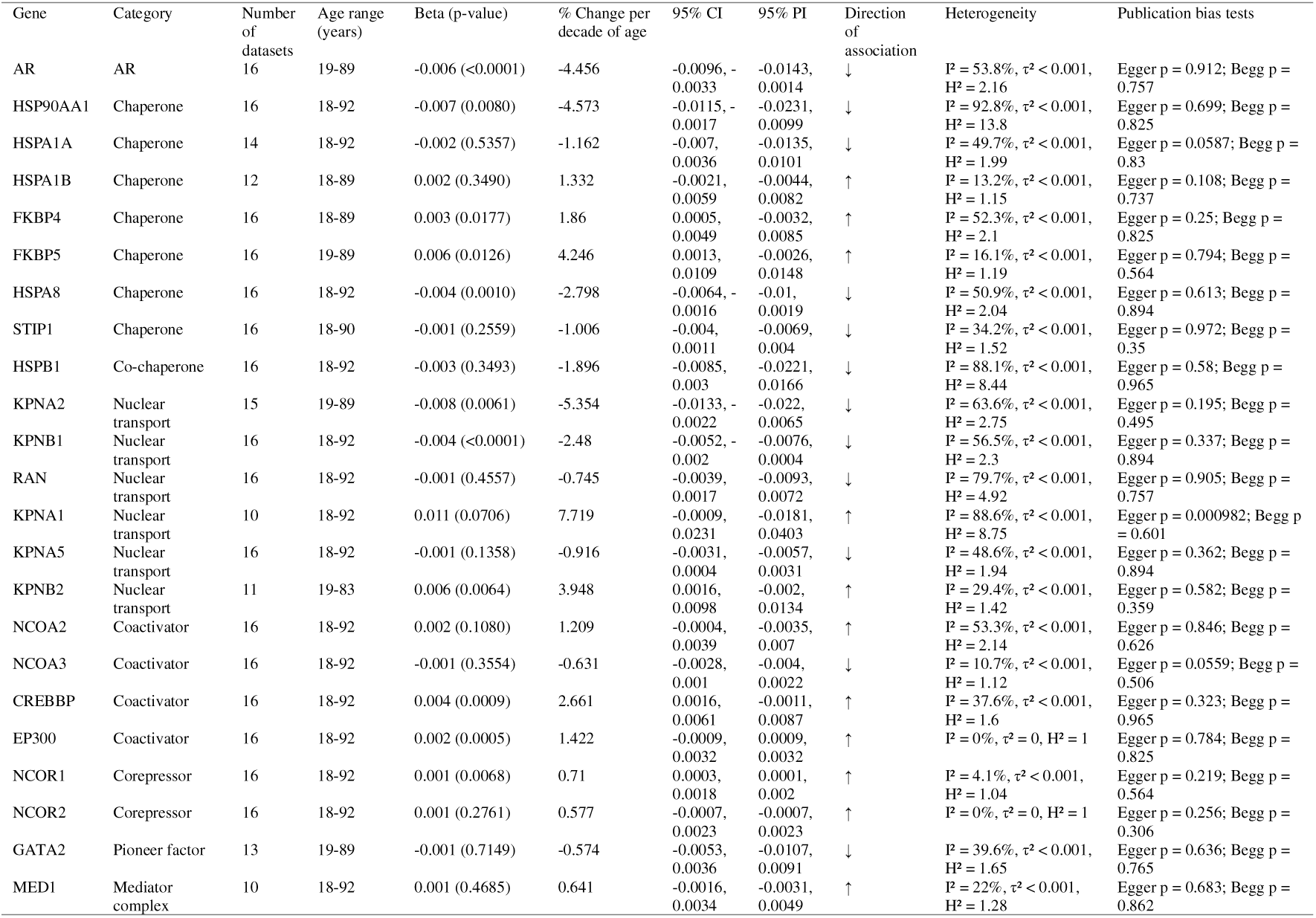
Gene-level meta-analysis results for association with age.

**Table 2.**
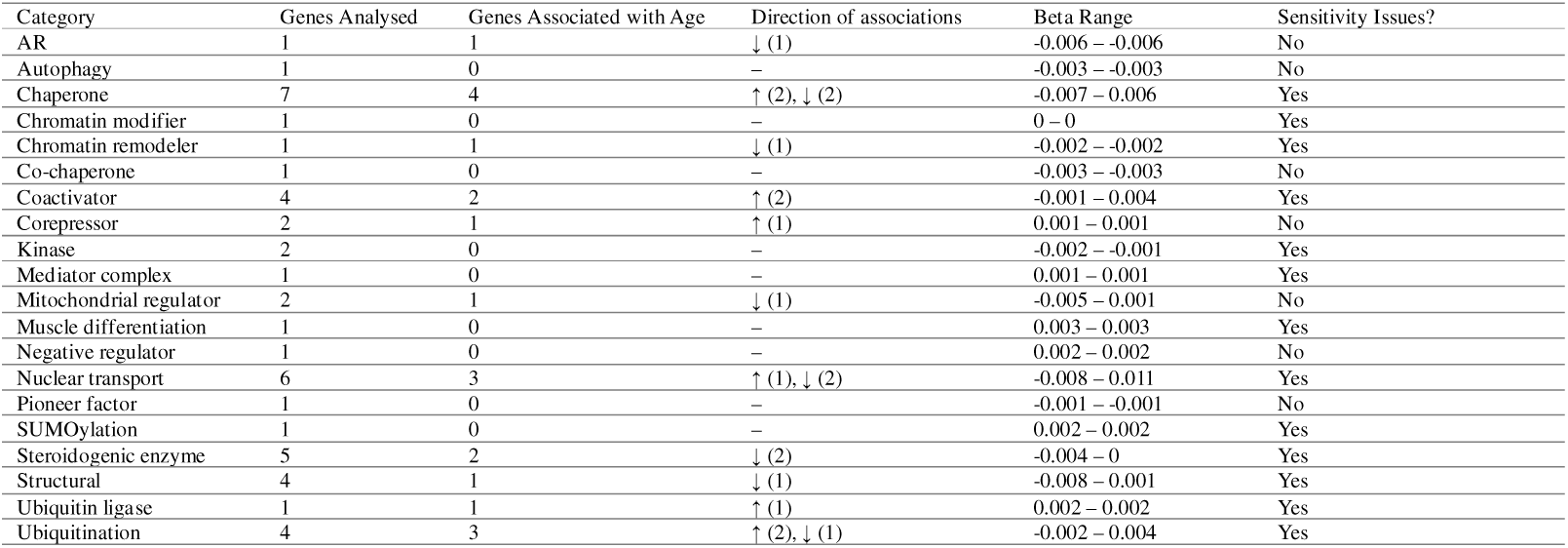
Summary of meta-analysis results by gene category. *LOO: Leave-one-out sensitivity analysis*

## Results

### Dataset characteristics

Systematic searching initially identified 618 datasets. Sixteen independent datasets were included in the final analysis comprising a total of 364 male skeletal muscle samples spanning an age range of 18-92 years (Figure 1). Muscle tissue was primarily sampled from the *vastus lateralis* (87% of datasets) and analysed using Illumina sequencing platforms (75% of datasets). Matrices were aligned to varying releases of the GRCh38, and in one case GRCh37, reference genome. The characteristics of each dataset are reported in full in Supplementary Table 1.

### Dataset quality appraisal

All 16 datasets met all critical and quality index criteria. One dataset did not specify the location of the muscle sample and instead defined the tissue as ‘skeletal muscle tissue near knee joint’. As we did not perform a meta-regression for tissue type, this was deemed acceptable for the purposes of this study. The results of the critical appraisal are presented in Supplementary Table 2.

### Meta analysis

#### AR expression is negatively associated with age

The relationship of age and AR expression was examined across 16 datasets (total n = 364, age range 18-92 years) (Supplementary Figure 1). A random effects meta-analysis indicated a significant negative association between AR expression and age (β = -0.006 log_2_ TMM-CPM per year, 95% CI: –0.0096 to –0.0033, *p* < 0.001) corresponding to a 4.4% decrease in expression per decade (Figure 2). There was moderate between-dataset heterogeneity (τ² = 0.000014, I² = 53.8%) (Supplementary Figure 2), and 95% prediction intervals (-0.0143 to 0.0014). Importantly, this indicates that future datasets were expected to show some variation in direction of effect. However, in this case, leave-one-out sensitivity analysis indicated that exclusion of any single dataset did not significantly affect the pooled effect estimate. There was no evidence of publication bias from visual inspection of funnel plots (Supplementary Figure 3), Egger’s regression test or Begg’s rank test (*p* > 0.05 for both).

**Figure 2.**
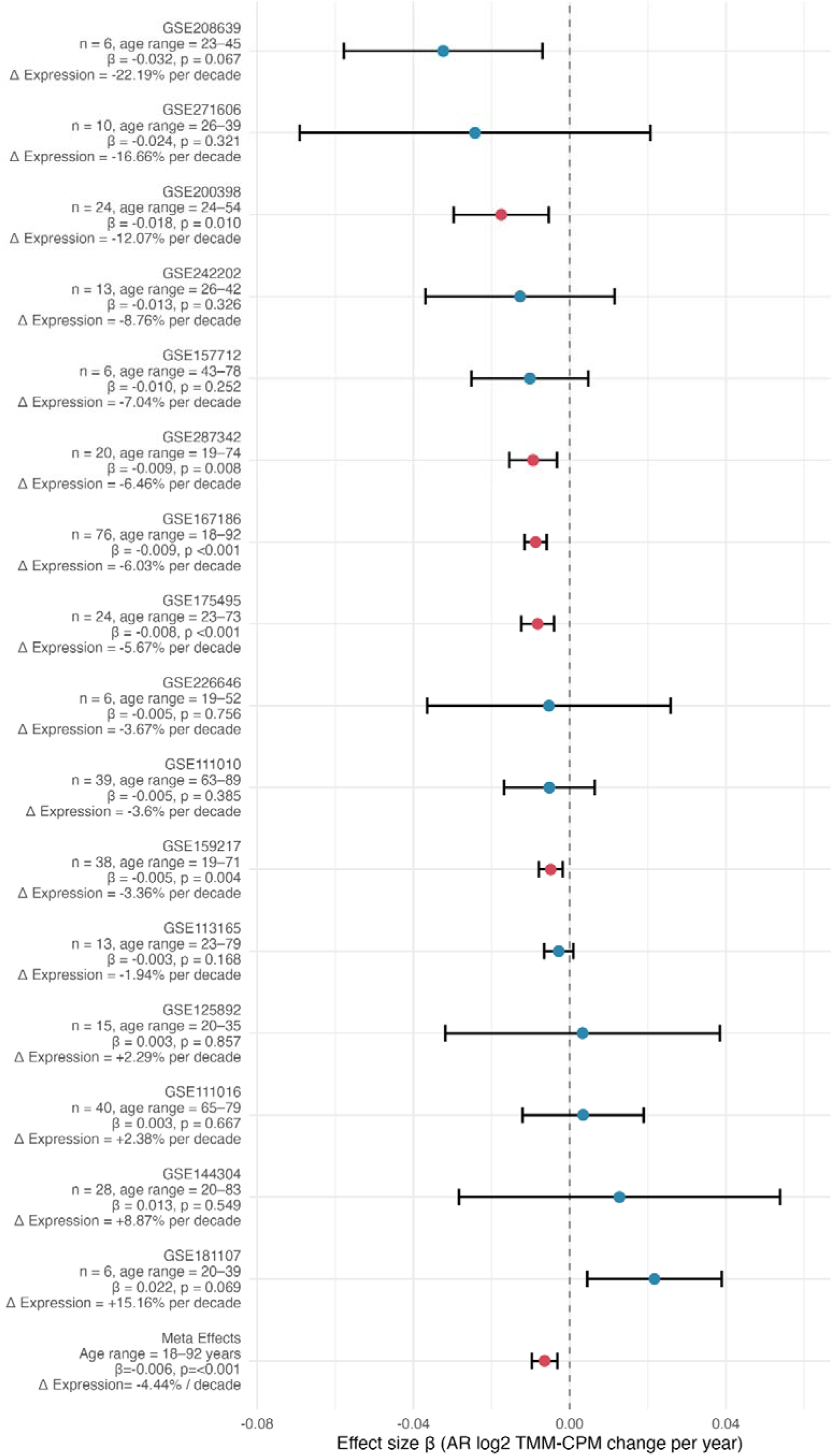
Forest plot of the skeletal muscle ARs association with age. *Point colour denotes statistical significance (red = p* < *0.05, blue = p* ≥ *0.05).* Δ *Expression denotes percentage change in expression of target gene per decade of age*.

### Age and AR transcriptional machinery

#### Transcriptional regulators

Of the seven chaperone genes investigated, two were positively associated with age (FKBP4, FKBP5; corresponding to a 1.86 and 4.24% increase in expression per decade respectively) and two were negatively associated with age (HSP90AA1; 4.57% decrease per decade, HSPA8; 2.79% decrease per decade) (Figure 3). HSPB1, a co-chaperone gene, showed no significant association with age. Of the four coactivator genes investigated, two were positively associated with age (CREBBP; 2.66% increase per decade, EP300; 1.42% increase per decade). Of the two corepressor genes investigated, one was positively associated with age (NCOR1; 0.71% increase per decade). The single chromatin remodeler investigated was negatively associated with age (SMARCA4; 1.2% decrease per decade). Neither of the two kinase genes, nor the single mediator complex, pioneer factor or chromatin modifier genes investigated, were significantly associated with age (Figure 3).

**Figure 3.**
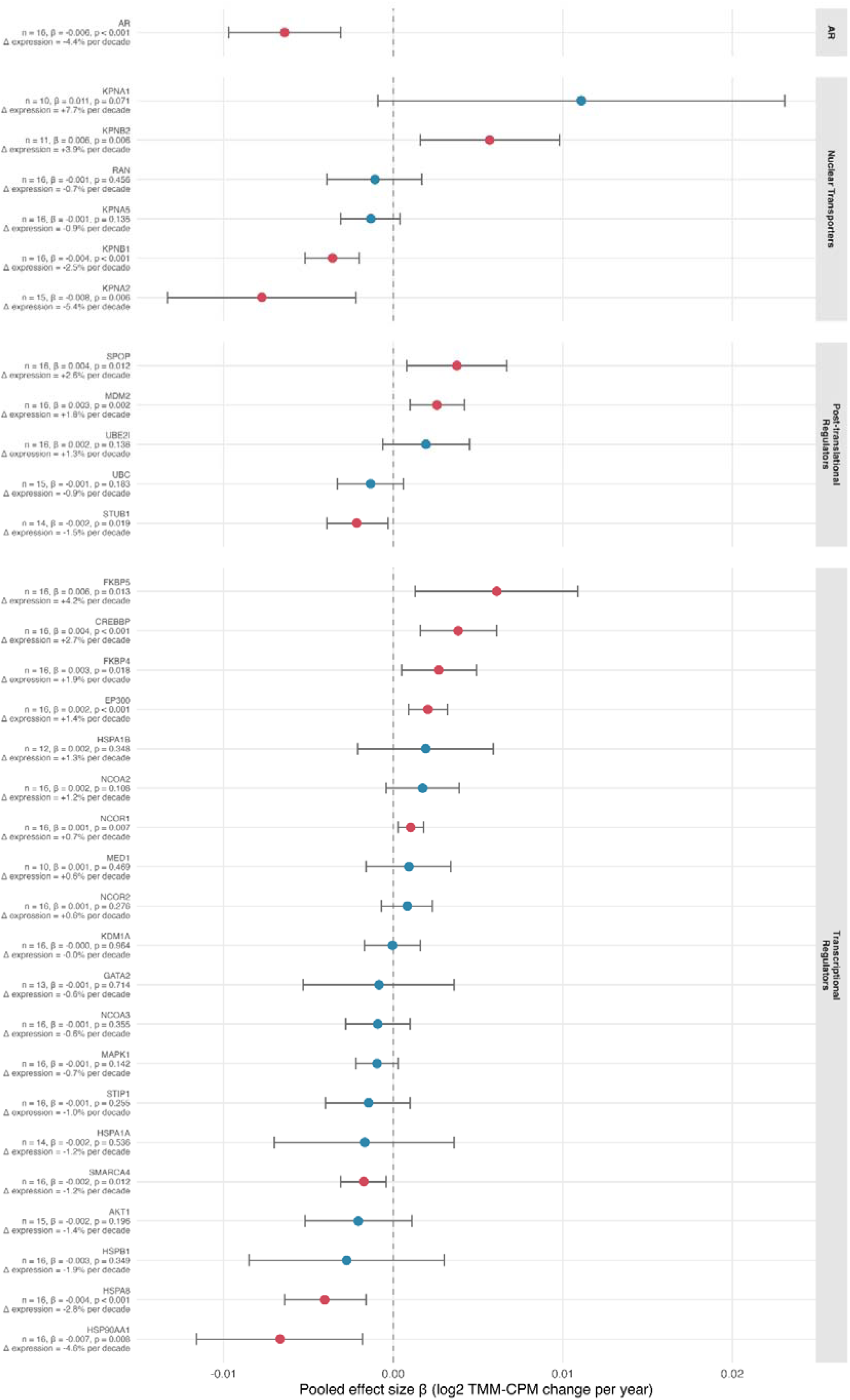
Forest plot of meta-analyses of AR-related genes association with age. *Point colour denotes statistical significance (red = p* < *0.05, blue = p* ≥ *0.05).* Δ *Expression denotes percentage change in expression of target gene per decade of age*.

#### Post-translational regulators

Of the five ubiquitination genes investigated, one was positively associated with age (MDM2; 1.78% increase per decade). The SUMOylation gene, UBE2I, was not significantly associated with age (Figure 3).

#### Nuclear transporters

Of the six nuclear transport genes, one was positively associated with age (KPNB2; 3.95% increase per decade) and two were negatively associated with age (KPNA2; 5.35% decrease per decade, KPNB1; 2.48% decrease per decade) (Figure 3).

#### Age and steroidogenic enzymes

Of the five steroidogenic enzymes investigated, two were negatively associated with age (STAR; 2.58% decrease per decade, HSD3B2; 1.69% decrease per decade) (Figure 4).

**Figure 4.**
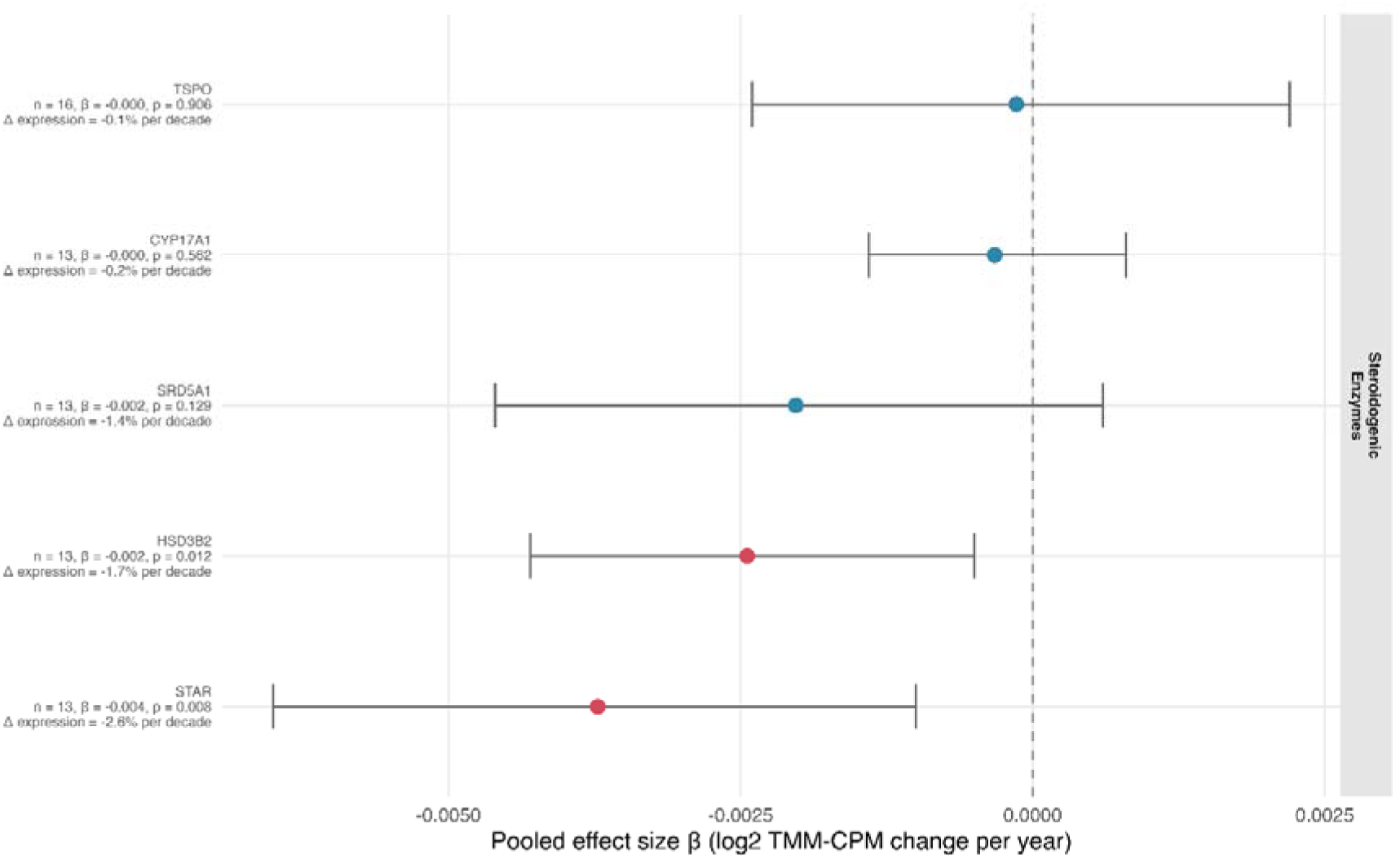
Forest plot of meta-analyses of steroidogenic genes association with age. *Point colour denotes statistical significance (red = p* < *0.05, blue = p* ≥ *0.05).* Δ *Expression denotes percentage change in expression of target gene per decade of age*.

#### Age and muscle signalling pathways

Of the four genes encoding structural muscle proteins, one was negatively associated with age (MYH2; 3.06% decrease per decade) (Figure 5). Of the two mitochondrial regulator genes investigated, one was negatively associated with age (PARGC1A; 3.79% decrease per decade) (figure 2.). The sole ubiquitin ligase gene was positively associated with age (TRIM63; 1.56% increase per decade) (Figure 5). Several genes involved in myogenic differentiation, autophagy and negative regulation of muscle mass (including MYOD1 and LC3B) were investigated but none were significantly associated with age (Figure 5).

**Figure 5.**
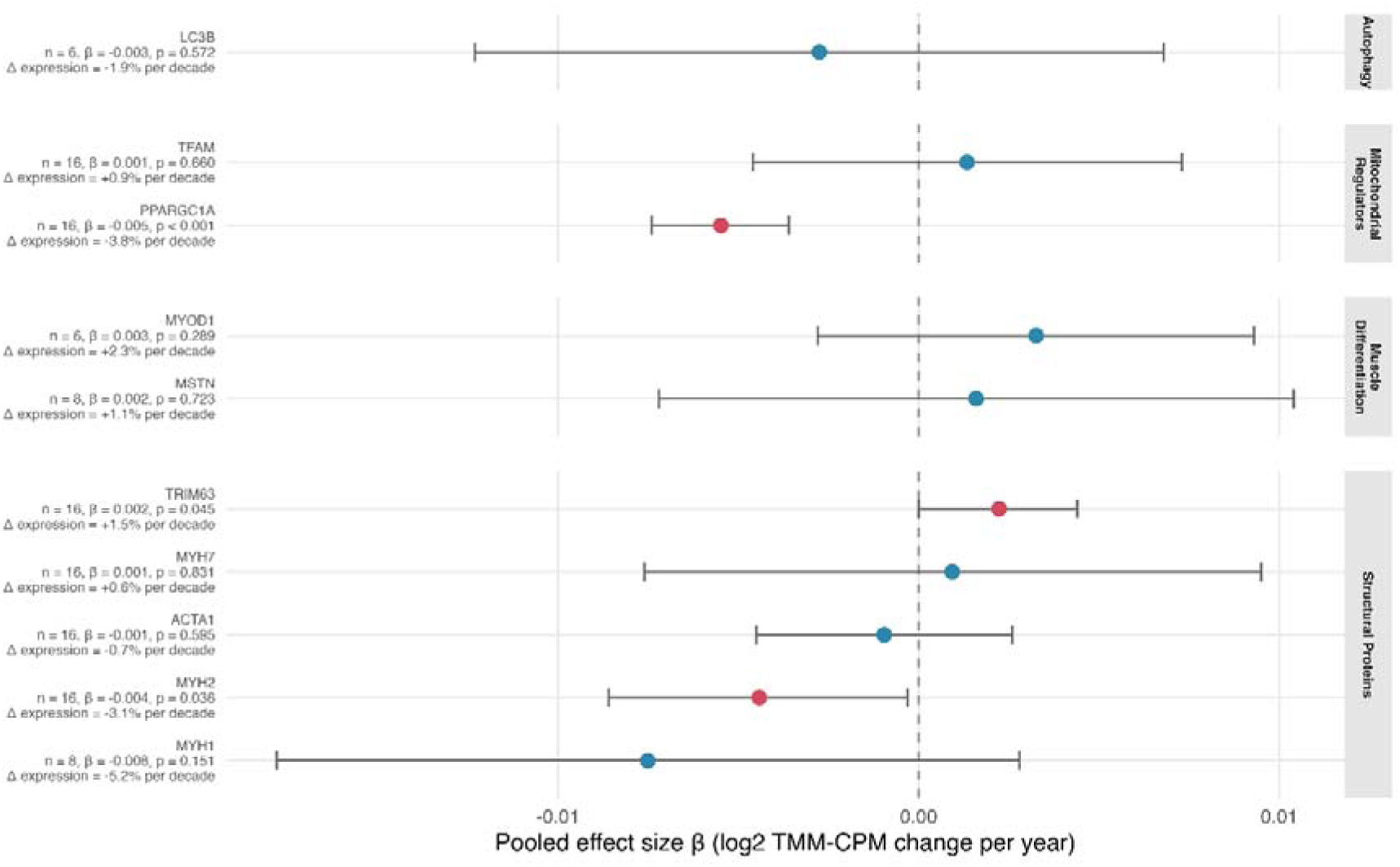
Forest plot of meta-analyses of AR target genes association with age. *Point colour denotes statistical significance (red = p* < *0.05, blue = p* ≥ *0.05).* Δ *Expression denotes percentage change in expression of target gene per decade of age*

## Discussion

The aim of this meta-analysis was to investigate the relationship between age and the expression levels of the AR gene, its associated transcriptional machinery and its downstream muscle signalling pathways. Our findings suggest that ageing in human skeletal muscle is associated with a targeted, multi-faceted remodeling of AR signalling, spanning a reduction in AR expression itself to changes in transcriptional regulation, nuclear transport and post-translational control of the AR (Figure 6).

**Figure 6.**
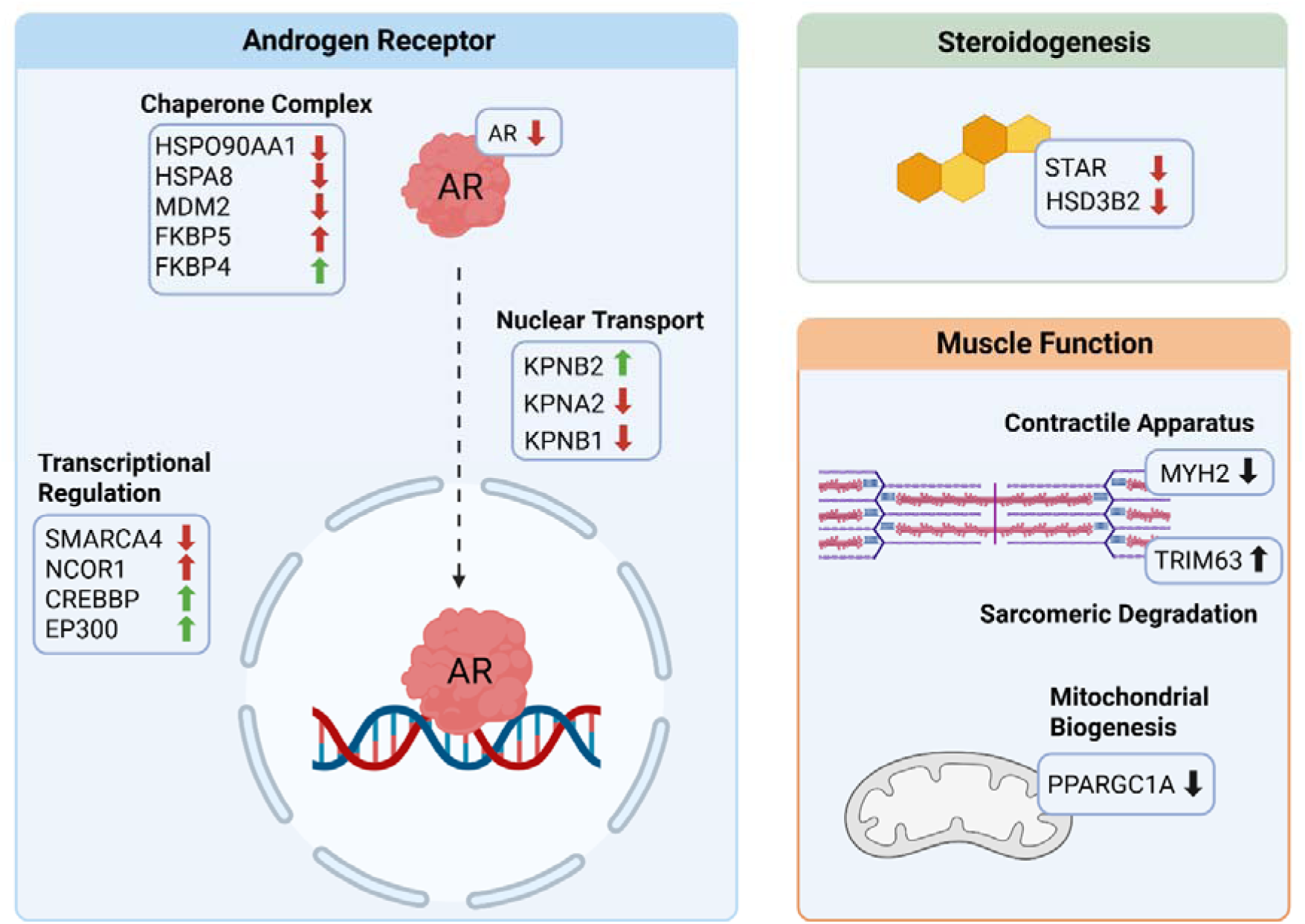
Effects of age on expression of AR-associated genes. *Arrow direction denotes effect of age on gene expression. Arrow colour denotes potential effect of change in gene expression on AR gene expression (Green = upregulated, Red = downregulated). Created with BioRender*.

### Ageing decreases AR expression, ligand binding and nuclear transport

Foremost, we report a reduction of 4.43% expression of the AR per year of age. It is however important to note that, despite the consistency of this observation, the same effect cannot be assumed across all future datasets. The moderate between-dataset heterogeneity (τ² = 0.000014, I² = 53.8%) and 95% prediction intervals (-0.0143 to 0.0014) crossing the zero line indicate that, even though leave-one-out sensitivity analysis indicated that exclusion of any single dataset did not significantly affect the pooled effect estimate in this case, the directional consistency of this effect cannot be assumed across all future datasets. In addition, it should be kept in mind that, due to the nature of our analysis, the reported percentage change per decade represents the overall age effect expressed on a per-decade scale for ease of interpretation, rather than a separately estimated effect for each decade of life.

A reduction in AR expression, and potentially consequent reduction in AR protein abundance, may therefore represent a bottleneck in androgenic signalling, potentially representing a compensatory mechanism that regulates the sensitivity of ageing skeletal muscle to physiological androgen exposure. We also observed an age-associated increase in MDM2 expression, which may promote greater proteasomal degradation of the receptor, actively reducing nuclear receptor content (56, 57), although the strength of this association showed some sensitivity to individual observations in the LOO analysis. In addition, the findings reported in this review are purely transcriptomic, and do not necessarily indicate an age-associated decline in AR protein abundance. Indeed, prior studies have reported no association between AR protein content and age in human skeletal muscle, albeit in cohorts of fewer than thirty participants that were primarily focused on the AR response to resistance training (41, 44, 58). However, negative associations with age have previously been reported in cross-sectional and longitudinal rat studies (59, 60). Nevertheless, studies are now required to confirm if AR protein abundance also declines with age in human male skeletal muscle consistent with our observations for gene expression.

Alongside a reduction in expression of the receptor itself, we observed negative associations between age and the chaperone genes HSP90AA1 and HSPA8, which may reduce AR protein stability and inhibit the targeting of misfolded proteins for degradation (61-63). Expression of FKBP5, a competitive binder to the HSP90-AR complex, was positively associated with age, potentially inhibiting nuclear translocation of the AR, however, LOO analysis suggested some variability in the robustness of this association (64, 65). Furthermore, we observed an age-associated decline in KPNA2 expression, suggesting further inhibition of classical nuclear import pathways and consequent cytoplasmic accumulation of the receptor (66-68). In contrast, KPNB2 expression was positively associated with age, possibly indicating an upregulation of alternative nuclear import channels. Therefore, it appears that as ageing may inhibit formation of the AR protein complex and close off classical nuclear import channels, alternative mechanisms for maintaining receptor complex binding and nuclear transport may be upregulated in response.

### Ageing is associated with changes in AR sensitivity

Several coactivator genes were positively associated with age, reflecting a potential compensatory, transcriptional response to diminishing receptor availability, with the aim of enhancing existing receptor signalling. Conversely, expression of the corepressor NCOR1 was positively associated with age, which may inhibit binding of transcriptional machinery and recruitment of said coactivators to the AR complex (69-71). The chromatin remodeler SMARCA4 was negatively associated with age, suggesting restricted chromatin access, and further repressing coactivator driven enhancement of AR transcription (72, 73), although the strength of this association showed some sensitivity to individual observations. Therefore, whilst AR coactivator genes appear to be upregulated with age, their influence on AR expression may be limited due to competition for binding sites with corepressors, and restricted chromatin access.

### Ageing is associated with selective transcriptional changes in steroidogenic enzyme expression, muscle fibre structure and mitochondrial regulation

To place our findings on AR signalling within the broader context of skeletal muscle ageing, we additionally examined associations between age and selected markers of muscle phenotype and metabolic function that are biologically relevant to, but not necessarily directly regulated by, androgen signalling. Upstream of the AR, we report a negative association of the steroidogenic enzymes STAR (albeit with significance sensitive to the inclusion of one dataset in leave-one-out analysis) and HSD3B2 with age, which may attenuate steroid hormone synthesis through a reduction in substrate availability (74-76). Among genes related to skeletal muscle phenotype that may be influenced by androgen signalling, expression of the fast type IIa muscle fibre encoding gene MYH2 but not type IIx encoding MYH1 or type I encoding MYH7, was negatively associated with age. Despite some dataset-level variability, this finding is consistent with evidence of preferential age-associated atrophy of the fast-compared with slow-twitch muscle fibre phenotype (77). Whilst prior histological studies suggest a preferential atrophy of type IIx rather than type IIa fibres (78, 79), our results indicate an age-associated reduction in MYH2 but not MYH1 expression. Although, our findings are derived from bulk transcriptomic data, in which extensive overlap of MYH1 and MYH2 expression occurs, preventing clear deconvolution of type IIa and type IIx fibres (80, 81). As such, whilst our results suggest selective transcriptional vulnerability of type IIa fibres, they should be interpreted with caution given the limitations of bulk RNA-seq.

Consistent with potential age-related changes in downstream androgen-responsive pathways, the mitochondrial regulator PARGC1A was negatively associated with age, indicating a reduction in metabolic and mitochondrial support of oxidative fibres. Such functional impairment of oxidative fibres is consistent with broader age-related declines in mitochondrial capacity and oxidative metabolism previously reported in human skeletal muscle (82-84).

Also consistent with prior research, various genes involved in muscle differentiation and anabolic growth signalling were not significantly associated with age (85-89). Collectively, the findings from *MetAR* suggest that ageing is primarily associated with downstream transcriptional changes related to skeletal muscle fibre structure and metabolic machinery, and not upstream anabolic and catabolic signalling.

### Strengths and limitations

A major component and strength of this study is the development and application of the *MetAR* pipeline. *MetAR* negates the requirement for high performance computing by acquiring counts matrices from NCBI GEO, thus allowing researchers to perform meta-analyses using standard computing resources. The pipeline provides significant flexibility in data handling, allowing researchers to tailor datasets to meet their specific inclusion criteria. Finally, whilst not included in this study, meta-regression analyses can also be integrated into the pipeline, to investigate various moderators such as sex or muscle type. These features make *MetAR* a versatile and accessible framework for transcriptomic analyses, with broad application across biological and biomedical research. Through the *MetAR* pipeline, we accounted for study level covariates such as disease status, acute exercise interventions and supplement/medication administration where these factors were reported as they may influence AR expression. However, this control was limited to information available in NCBI GEO metadata and the participant inclusion criteria reported in the corresponding publications. As such, we could not implement consistent controls for other potential confounders, including habitual exercise, diet or exposure to medications that may affect AR expression (41, 90, 91). These factors are important moderators of mRNA expression, and future implementations of the *MetAR* pipeline may go to more extensive lengths to retrieve additional data from custodians, allowing for extensive control of covariates. As reported in supplementary table 1, although several datasets incorporated broad age ranges of participants, many sampled individuals from the extreme ends of the ageing spectrum, typically comparing groups of young and older adult males. In addition, as noted above, the reported percentage change per decade reflects the overall age effect expressed on a per-decade scale. Consequently, our analysis is limited in its ability to describe more dynamic changes in AR expression across the male lifespan. The most prominent limitation of this study is its purely transcriptomic nature, with no integrated proteomic validation to confirm whether observed associations between gene expression and age are reflected in protein abundance. Furthermore, effect sizes for many reported associations were small, and subtle biological effects may therefore be obscured. There was some variability across datasets in terms of sequencing platform, alignment method and libraries. Conversely, there was minimal heterogeneity in the specific muscle sampled. Prior longitudinal rodent work suggests a muscle type-specific response of the AR to ageing at the protein level, whereby AR protein declined in the predominantly oxidative soleus muscle, but not in the glycolytic plantaris (59). AR protein is also more abundant in young rodents in predominantly oxidative muscle, as such, muscle phenotypes more abundant in AR content may be more susceptible, or the first to exhibit, age-associated reductions in AR expression (92, 93). Collectively, these findings do not establish a mechanistic link but do suggest that the oxidative muscle phenotype itself may be vulnerable to age-associated changes in the AR, which could be explored in future meta-regression analyses.

Finally, while the current study focuses on males only, recent evidence from our group suggests that androgen availability and AR activation are associated with skeletal muscle phenotypes in females (94). Whether skeletal muscle AR expression and its regulatory machinery undergo similar age-associated changes in females however remains unexplored. Future studies directly comparing males and females will therefore be important to determine whether the molecular changes identified in this study represent conserved or sex-specific features of skeletal muscle ageing.

### Conclusion

Our findings suggest that ageing is associated with a reduction in skeletal muscle AR expression, alongside a decreased expression of genes promoting AR stability, ligand binding and nuclear transport. These changes are accompanied by some compensatory mechanisms, including transcriptional upregulation of AR coactivators and alternative nuclear transport channels. Ageing was linked to transcriptional changes affecting muscle fibre structure and metabolic capacity, primarily in genes associated with the fast-twitch phenotype. Conversely, genes associated with anabolic growth signalling and differentiation in muscle remained stable with age, consistent with prior research.

## Additional Information

### Contributors

- Ross Williams contributed to study concept and design, data collection, data analysis, drafting and critical revision of the paper.
- Viktor Engman contributed to data collection, drafting and critical revision of the paper.
- Danielle Hiam contributed to data analysis, drafting and critical revision of the paper.
- Megan Soria contributed to data analysis and critical revision of the paper.
- Glenn Wadley contributed to drafting and critical revision of the paper.
- Séverine Lamon contributed to funding the study, drafting and critical revision of the paper.
- All authors reviewed and approved the final version, and no other person made a substantial contribution to the paper.

### Ethical standards

All authors certify that they comply with the ethical guidelines for authorship and publishing in Physiological Genomics. The manuscript does not contain primary clinical studies or identifiable patient data.

### Supplementary materials

- Supplementary Figure 1 – Dataset-specific associations between AR gene expression and age
- Supplementary Figure 2 – Baujat plot for the meta-analysis of the association between AR gene expression and age
- Supplementary Figure 3 – Funnel plot assessing publication bias for the meta-analysis of the association between AR gene expression and age
- Supplementary Table 1 – Characteristics of included datasets
- Supplementary Table 2 – Quality appraisal of included datasets
- Supplementary Table 3 – Leave-one-out analysis
- Supplementary Table 4 – Leave-one-out analysis: Summary of influential datasets

