## Supplementary Figures for "*MetAR*: A semi-automated meta-analysis of skeletal muscle androgen receptors association with age"

Supplementary Figure 1. Dataset-specific associations between AR gene expression and age. *Scatterplots show the relationship between age and AR expression for each dataset included in the meta-analysis. Fitted regression lines indicate direction and magnitude of association. Shaded regions indicate 95% confidence intervals.*

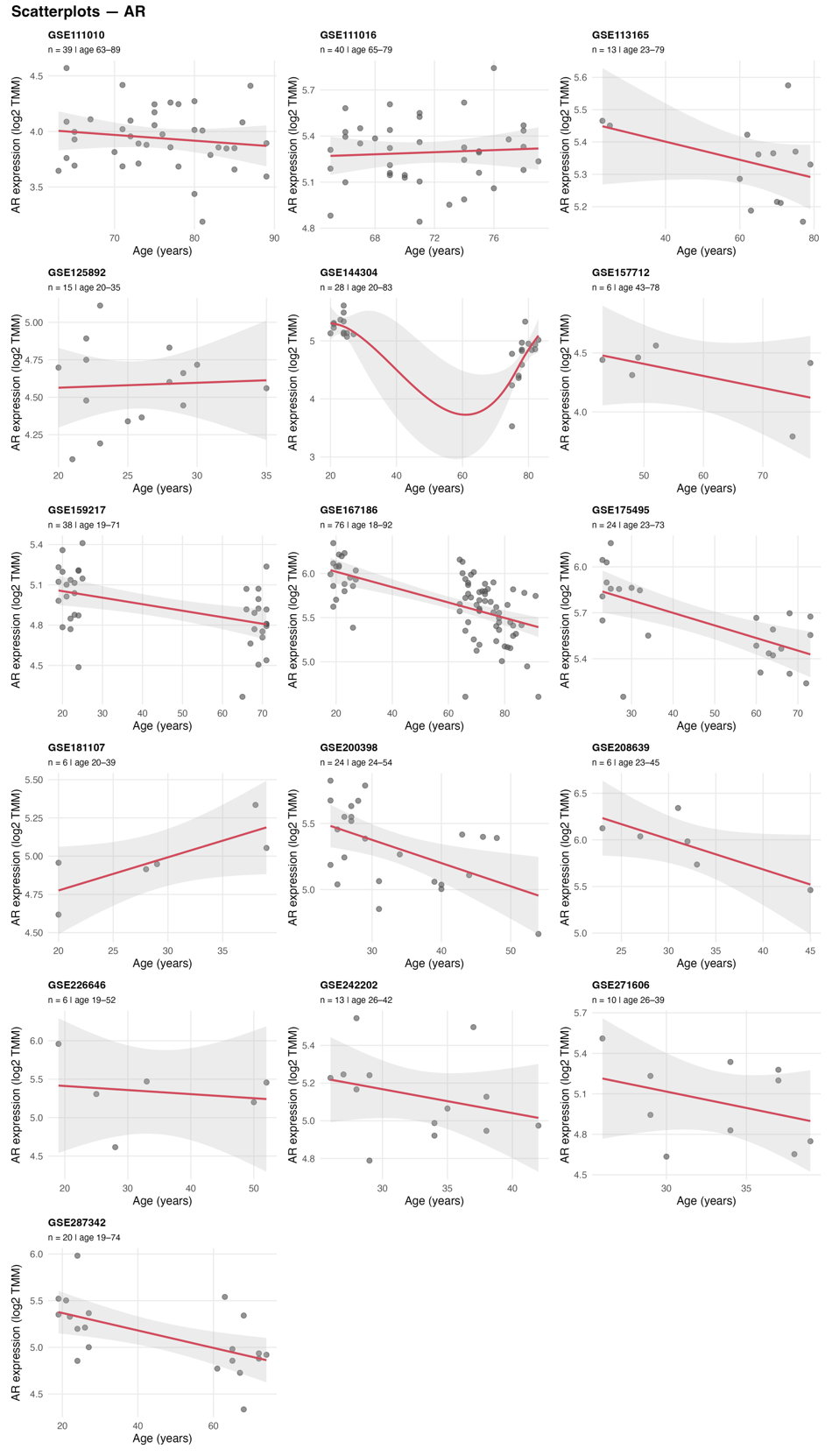

Supplementary Figure 2. Baujat plot for the meta-analysis of the association between AR gene expression and age. *The contribution of each dataset to overall heterogeneity (Cochran’s Q) is plotted against its influence on the pooled effect size.*

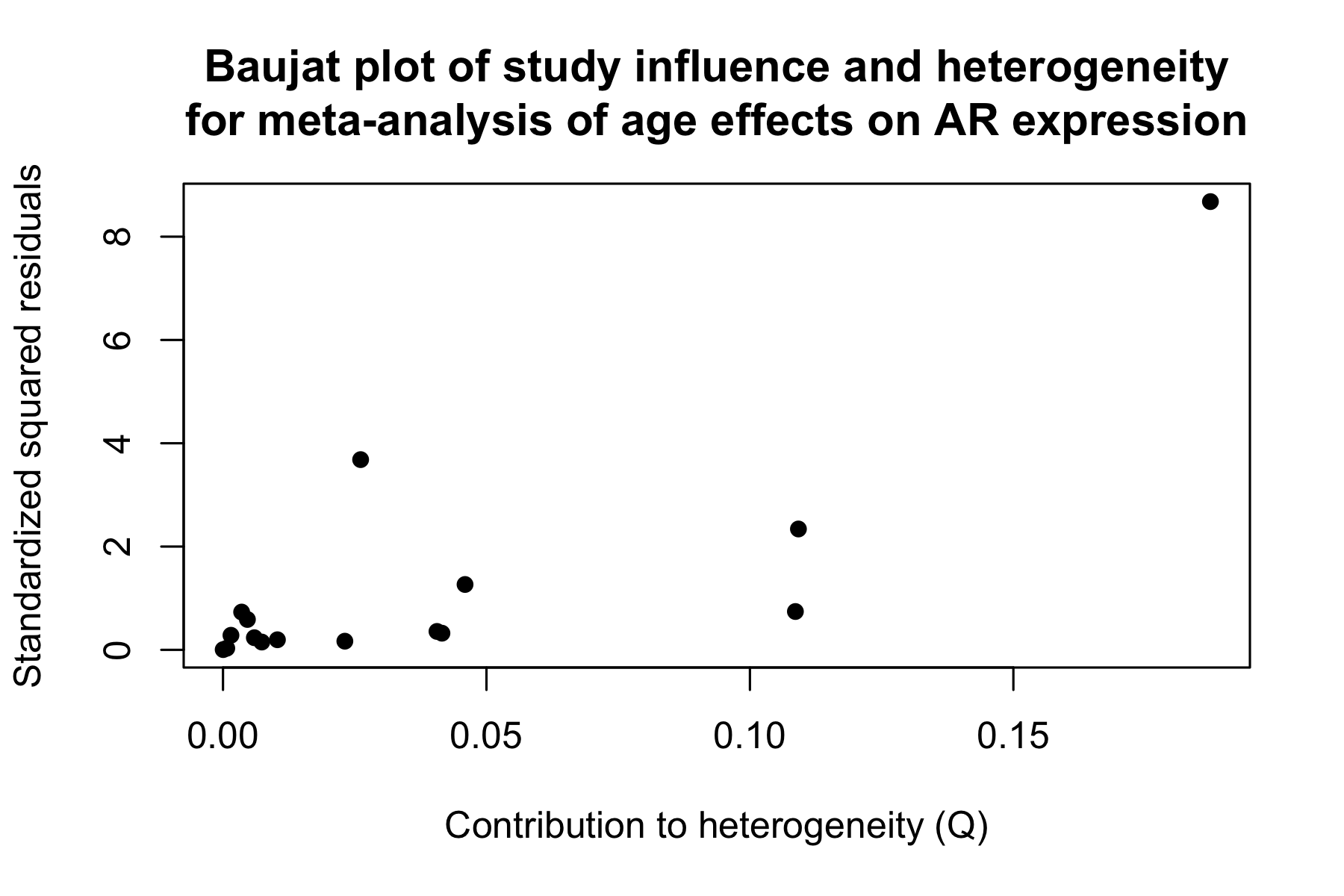

Supplementary Figure 3. Funnel Plot assessing publication bias for the meta-analysis of the association between AR gene expression and age.

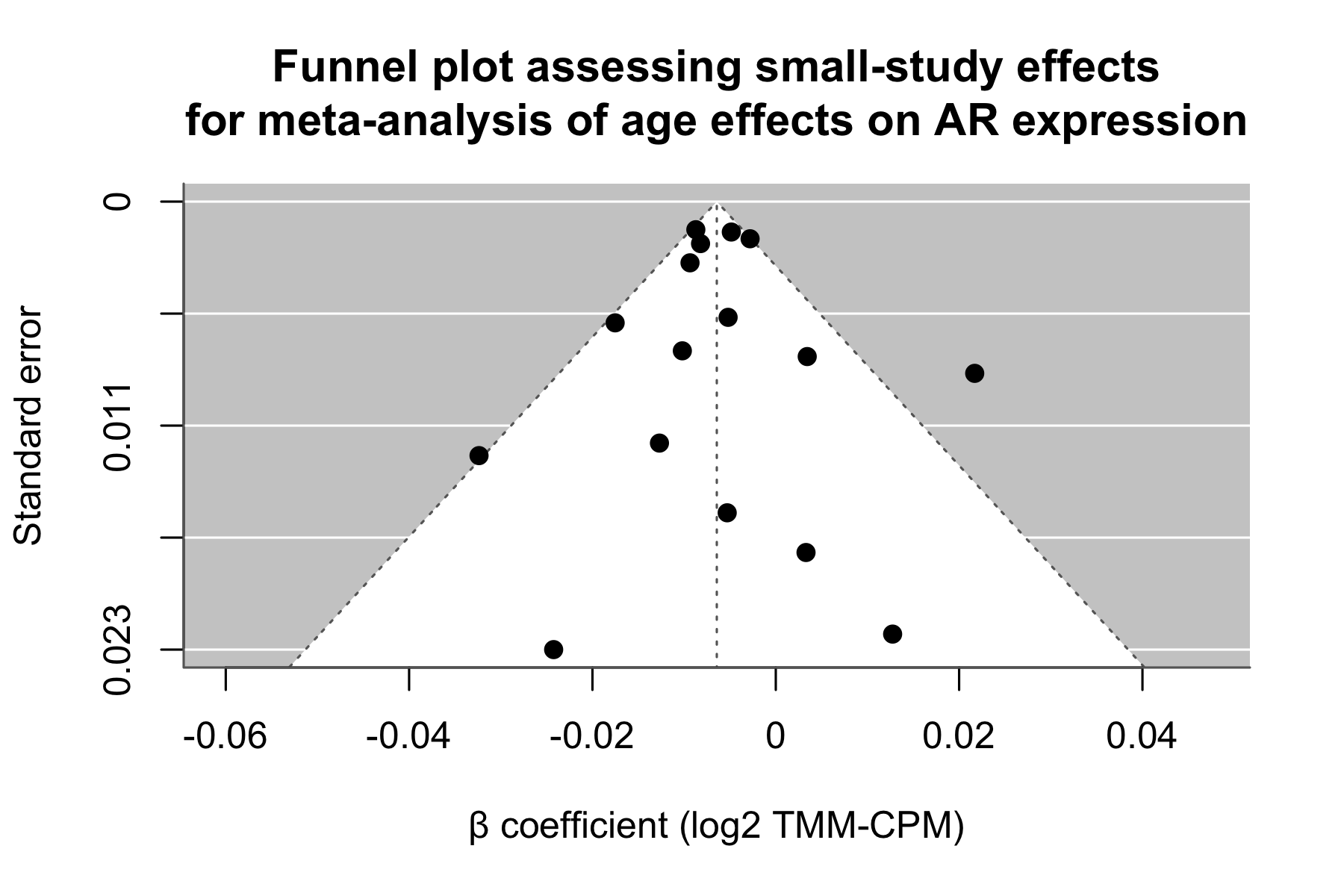
