## Supplementary Tables for "*MetAR*: A semi-automated meta-analysis of skeletal muscle androgen receptors association with age"

Supplementary Table 1. Characteristics of included datasets

| Dataset Accession ID | Sample Size | Mean age (years) ± SD (Range) | Tissue | Sequencing Instrument | Platform ID | Reference Genome | Library Preparation Method | Sequencing depth (M) | Alignment Method |
| --- | --- | --- | --- | --- | --- | --- | --- | --- | --- |
| GSE111010 | 39 | 75.6 ± 7.6 (63.0–89.0) | Vastus lateralis | Illumina HiSeq 2500 | GPL16791 | GRCh38.p2 | TruSeq Stranded RNA Ribo Zero Gold Kit (Illumina) | 34-110 | STAR |
| GSE111016 | 40 | 71.5 ± 4.3 (65.0–79.0) | Vastus lateralis | Illumina HiSeq 2500 | GPL16791 | GRCh38.p2 | TruSeq Stranded RNA Ribo Zero Gold Kit (Illumina) | 77.3 | STAR |
| GSE113165 | 13 | 62.5 ± 18.0 (23.0–79.0) | Vastus lateralis | Illumina HiSeq 2500 | GPL16791 | GRCh38 | TruSeq Stranded RNA Ribo Zero Gold Kit (Illumina) | 77.3 | STAR |
| GSE125892 | 15 | 25.5 ± 4.2 (20.0–35.0) | Vastus lateralis | AB 5500xl Genetic Analyzer | GPL16288 | GRCh38 | Applied Biosystems SOLiDSAGE kit (SOLiD 4 System) | 13.9 | SOLiD SAGE |
| GSE144304 | 28 | 52.9 ± 28.2 (20.4–82.7) | Vastus lateralis | Illumina NextSeq 500 | GPL18573 | GRCh38 | NEBNext Ultra Directional RNA Library Prep Kit | 16.4 | STAR |
| GSE157712 | 6 | 57.5 ± 15.0 (43.0–78.0) | Vastus lateralis | Illumina HiSeq 4000 | GPL20301 | GRCh38 | NEBNext Ultra II Directional RNA Library Prep Kit with NEBNext rRNA Depletion Module | 29.7 | STAR |
| GSE159217 | 38 | 44.3 ± 23.9 (19.0–71.0) | Vastus lateralis | BGISEQ-500 | GPL23227 | GRCh38.p13 | Not described - standard BGISEQ-500 library prep | 41.6 | STAR |
| GSE167186 | 76 | 61.2 ± 23.7 (18.0–92.0) | Vastus lateralis | Illumina HiSeq 4000 | GPL20301 | GRCh38 | Batch-tag-seq libraries (BGI-style) | 3.5 | STAR |
| GSE175495 | 24 | 46.2 ± 20.6 (23.0–73.0) | Vastus lateralis | Illumina HiSeq 4000 | GPL20301 | GRCh38.p12 | NEBNext Ultra II mRNA kit (RiboZero-depleted total RNA) | 51.3 | HISAT2 |
| GSE181107 | 6 | 29.0 ± 8.3 (20.0–39.0) | Vastus lateralis* | BGISEQ-500 | GPL23227 | GRCh38.p13 | PolyA-selected cDNA | 44.5 | STAR |
| GSE200398 | 24 | 33.0 ± 8.9 (24.0–54.0) | Vastus lateralis | Illumina HiSeq 2500 | GPL16791 | GRCh37 | NEBNext Ultra II Directional RNA Library Prep Kit with NEBNext rRNA Depletion Module | 40.5 | Kallisto |
| GSE208639 | 6 | 31.8 ± 7.4 (23.0–45.0) | Vastus lateralis | Illumina NextSeq 500 | GPL18573 | GRCh38 | NEBNext Ultra II Directional RNA Library Prep Kit with NEBNext rRNA Depletion Module | 64.8 | STAR |
| GSE226646 | 6 | 34.5 ± 13.6 (19.0–52.0) | Gastrocnemius | Ion Torrent Proton | GPL17303 | GRCh38 | Lexogen QuantSeq 3′ mRNA Kit | 6 | STAR |
| GSE242202 | 13 | 32.7 ± 5.2 (26.0–42.0) | Vastus lateralis | Illumina NextSeq 550 | GPL21697 | GRCh38.p13 | NEBNext Ultra II Directional RNA Library Prep Kit | 64.8 | HISAT2 v2.1.0 |
| GSE271606 | 10 | 33.3 ± 4.5 (26.0–39.0) | Vastus lateralis | Illumina NextSeq 550 | GPL21697 | GRCh38.p13 | NEBNext Ultra II Directional RNA Library Prep Kit | 59.4 | HISAT2 v2.1.0 |
| GSE287342 | 20 | 45.4 ± 22.9 (19.0–74.0) | Vastus lateralis | Illumina NovaSeq 6000 | GPL24676 | GRCh38.p14 | TruSeq Stranded Total RNA Library Prep Gold Kit (Ribo-Zero Gold) | 34 | Kallisto |

Supplementary Table 2. Quality appraisal of included datasets. **Defined only as ‘skeletal muscle tissue near knee joint’ – most likely vastus lateralis*

| *Critical Criteria* | | | | | | |
| --- | --- | --- | --- | --- | --- | --- |
| Dataset Accession ID | Age defined? | Sex defined? | Tissue defined? | Raw counts available? | Basal gene expression |  |
| GSE111010 | ✓ | ✓ | ✓ | ✓ | ✓ |  |
| GSE111016 | ✓ | ✓ | ✓ | ✓ | ✓ |  |
| GSE113165 | ✓ | ✓ | ✓ | ✓ | ✓ |  |
| GSE125892 | ✓ | ✓ | ✓ | ✓ | ✓ |  |
| GSE144304 | ✓ | ✓ | ✓ | ✓ | ✓ |  |
| GSE157712 | ✓ | ✓ | ✓ | ✓ | ✓ |  |
| GSE159217 | ✓ | ✓ | ✓ | ✓ | ✓ |  |
| GSE167186 | ✓ | ✓ | ✓ | ✓ | ✓ |  |
| GSE175495 | ✓ | ✓ | ✓ | ✓ | ✓ |  |
| GSE181107 | ✓ | ✓ | ✓* | ✓ | ✓ |  |
| GSE200398 | ✓ | ✓ | ✓ | ✓ | ✓ |  |
| GSE208639 | ✓ | ✓ | ✓ | ✓ | ✓ |  |
| GSE226646 | ✓ | ✓ | ✓ | ✓ | ✓ |  |
| GSE242202 | ✓ | ✓ | ✓ | ✓ | ✓ |  |
| GSE271606 | ✓ | ✓ | ✓ | ✓ | ✓ |  |
| GSE287342 | ✓ | ✓ | ✓ | ✓ | ✓ |  |
| *Quality Assessment* | | | | | | |
| Dataset Accession ID | Sequencing platform defined? | Library preparation defined? | Alignment method defined? | Genome build defined? | Counts generation method defined? | Sample ID’s match metadata? |
| GSE111010 | ✓ | ✓ | ✓ | ✓ | ✓ | ✓ |
| GSE111016 | ✓ | ✓ | ✓ | ✓ | ✓ | ✓ |
| GSE113165 | ✓ | ✓ | ✓ | ✓ | ✓ | ✓ |
| GSE125892 | ✓ | ✓ | ✓ | ✓ | ✓ | ✓ |
| GSE144304 | ✓ | ✓ | ✓ | ✓ | ✓ | ✓ |
| GSE157712 | ✓ | ✓ | ✓ | ✓ | ✓ | ✓ |
| GSE159217 | ✓ | ✓ | ✓ | ✓ | ✓ | ✓ |
| GSE167186 | ✓ | ✓ | ✓ | ✓ | ✓ | ✓ |
| GSE175495 | ✓ | ✓ | ✓ | ✓ | ✓ | ✓ |
| GSE181107 | ✓ | ✓ | ✓ | ✓ | ✓ | ✓ |
| GSE200398 | ✓ | ✓ | ✓ | ✓ | ✓ | ✓ |
| GSE208639 | ✓ | ✓ | ✓ | ✓ | ✓ | ✓ |
| GSE226646 | ✓ | ✓ | ✓ | ✓ | ✓ | ✓ |
| GSE242202 | ✓ | ✓ | ✓ | ✓ | ✓ | ✓ |
| GSE271606 | ✓ | ✓ | ✓ | ✓ | ✓ | ✓ |
| GSE287342 | ✓ | ✓ | ✓ | ✓ | ✓ | ✓ |

Supplementary table 3. Leave-one-out analysis. *LOO: Leave-one-out*

| Gene | Category | Excluded Dataset | Full Model Beta (p-value) | LOO Model Beta (p-value) | Impact of dataset exclusion |
| --- | --- | --- | --- | --- | --- |
| AR | AR | GSE111010 | -0.007 (<0.0001) | -0.007 (<0.0001) | No meaningful change |
| AR | AR | GSE111016 | -0.007 (<0.0001) | -0.007 (<0.0001) | No meaningful change |
| AR | AR | GSE113165 | -0.007 (<0.0001) | -0.007 (<0.0001) | No meaningful change |
| AR | AR | GSE125892 | -0.007 (<0.0001) | -0.007 (<0.0001) | No meaningful change |
| AR | AR | GSE144304 | -0.007 (<0.0001) | -0.007 (0.0001) | No meaningful change |
| AR | AR | GSE157712 | -0.007 (<0.0001) | -0.006 (<0.0001) | No meaningful change |
| AR | AR | GSE159217 | -0.007 (<0.0001) | -0.007 (<0.0001) | No meaningful change |
| AR | AR | GSE167186 | -0.007 (<0.0001) | -0.006 (<0.0001) | No meaningful change |
| AR | AR | GSE175495 | -0.007 (<0.0001) | -0.006 (<0.0001) | No meaningful change |
| AR | AR | GSE181107 | -0.007 (<0.0001) | -0.007 (<0.0001) | No meaningful change |
| AR | AR | GSE200398 | -0.007 (<0.0001) | -0.006 (<0.0001) | No meaningful change |
| AR | AR | GSE208639 | -0.007 (<0.0001) | -0.006 (<0.0001) | No meaningful change |
| AR | AR | GSE226646 | -0.007 (<0.0001) | -0.007 (<0.0001) | No meaningful change |
| AR | AR | GSE242202 | -0.007 (<0.0001) | -0.006 (<0.0001) | No meaningful change |
| AR | AR | GSE271606 | -0.007 (<0.0001) | -0.006 (<0.0001) | No meaningful change |
| AR | AR | GSE287342 | -0.007 (<0.0001) | -0.006 (<0.0001) | No meaningful change |
| HSP90AA1 | Chaperone | GSE111010 | -0.007 (0.0068) | -0.007 (0.0082) | No meaningful change |
| HSP90AA1 | Chaperone | GSE111016 | -0.007 (0.0068) | -0.007 (0.0165) | No meaningful change |
| HSP90AA1 | Chaperone | GSE113165 | -0.007 (0.0068) | -0.006 (0.0186) | No meaningful change |
| HSP90AA1 | Chaperone | GSE125892 | -0.007 (0.0068) | -0.007 (0.0115) | No meaningful change |
| HSP90AA1 | Chaperone | GSE144304 | -0.007 (0.0068) | -0.008 (0.0057) | No meaningful change |
| HSP90AA1 | Chaperone | GSE157712 | -0.007 (0.0068) | -0.007 (0.0103) | No meaningful change |
| HSP90AA1 | Chaperone | GSE159217 | -0.007 (0.0068) | -0.007 (0.0109) | No meaningful change |
| HSP90AA1 | Chaperone | GSE167186 | -0.007 (0.0068) | -0.008 (0.0058) | No meaningful change |
| HSP90AA1 | Chaperone | GSE175495 | -0.007 (0.0068) | -0.008 (0.0057) | No meaningful change |
| HSP90AA1 | Chaperone | GSE181107 | -0.007 (0.0068) | -0.007 (0.0054) | No meaningful change |
| HSP90AA1 | Chaperone | GSE200398 | -0.007 (0.0068) | -0.008 (0.0060) | No meaningful change |
| HSP90AA1 | Chaperone | GSE208639 | -0.007 (0.0068) | -0.004 (0.0047) | No meaningful change |
| HSP90AA1 | Chaperone | GSE226646 | -0.007 (0.0068) | -0.007 (0.0078) | No meaningful change |
| HSP90AA1 | Chaperone | GSE242202 | -0.007 (0.0068) | -0.008 (0.0054) | No meaningful change |
| HSP90AA1 | Chaperone | GSE271606 | -0.007 (0.0068) | -0.007 (0.0084) | No meaningful change |
| HSP90AA1 | Chaperone | GSE287342 | -0.007 (0.0068) | -0.007 (0.0123) | No meaningful change |
| HSPA1A | Chaperone | GSE111010 | -0.001 (0.6106) | -0.002 (0.5603) | No meaningful change |
| HSPA1A | Chaperone | GSE111016 | -0.001 (0.6106) | -0.001 (0.6231) | No meaningful change |
| HSPA1A | Chaperone | GSE113165 | -0.001 (0.6106) | 0.000 (0.9049) | No meaningful change |
| HSPA1A | Chaperone | GSE125892 | -0.001 (0.6106) | -0.002 (0.5757) | No meaningful change |
| HSPA1A | Chaperone | GSE144304 | -0.001 (0.6106) | -0.003 (0.1878) | No meaningful change |
| HSPA1A | Chaperone | GSE157712 | -0.001 (0.6106) | -0.001 (0.6157) | No meaningful change |
| HSPA1A | Chaperone | GSE159217 | -0.001 (0.6106) | -0.002 (0.6318) | No meaningful change |
| HSPA1A | Chaperone | GSE167186 | -0.001 (0.6106) | -0.003 (0.3984) | No meaningful change |
| HSPA1A | Chaperone | GSE181107 | -0.001 (0.6106) | -0.001 (0.6102) | No meaningful change |
| HSPA1A | Chaperone | GSE200398 | -0.001 (0.6106) | -0.001 (0.6928) | No meaningful change |
| HSPA1A | Chaperone | GSE208639 | -0.001 (0.6106) | -0.001 (0.6359) | No meaningful change |
| HSPA1A | Chaperone | GSE226646 | -0.001 (0.6106) | -0.001 (0.7700) | No meaningful change |
| HSPA1A | Chaperone | GSE242202 | -0.001 (0.6106) | -0.001 (0.5986) | No meaningful change |
| HSPA1A | Chaperone | GSE271606 | -0.001 (0.6106) | -0.001 (0.7224) | No meaningful change |
| HSPA1B | Chaperone | GSE111010 | 0.002 (0.3627) | 0.002 (0.5044) | No meaningful change |
| HSPA1B | Chaperone | GSE111016 | 0.002 (0.3627) | 0.002 (0.3525) | No meaningful change |
| HSPA1B | Chaperone | GSE113165 | 0.002 (0.3627) | 0.002 (0.3252) | No meaningful change |
| HSPA1B | Chaperone | GSE144304 | 0.002 (0.3627) | 0.000 (0.9602) | No meaningful change |
| HSPA1B | Chaperone | GSE157712 | 0.002 (0.3627) | 0.002 (0.3025) | No meaningful change |
| HSPA1B | Chaperone | GSE159217 | 0.002 (0.3627) | 0.002 (0.5692) | No meaningful change |
| HSPA1B | Chaperone | GSE181107 | 0.002 (0.3627) | 0.002 (0.3508) | No meaningful change |
| HSPA1B | Chaperone | GSE200398 | 0.002 (0.3627) | 0.003 (0.2154) | No meaningful change |
| HSPA1B | Chaperone | GSE208639 | 0.002 (0.3627) | 0.002 (0.3407) | No meaningful change |
| HSPA1B | Chaperone | GSE226646 | 0.002 (0.3627) | 0.002 (0.3424) | No meaningful change |
| HSPA1B | Chaperone | GSE242202 | 0.002 (0.3627) | 0.002 (0.3921) | No meaningful change |
| HSPA1B | Chaperone | GSE271606 | 0.002 (0.3627) | 0.002 (0.2965) | No meaningful change |
| FKBP4 | Chaperone | GSE111010 | 0.003 (0.0177) | 0.002 (0.0543) | Significance changed |
| FKBP4 | Chaperone | GSE111016 | 0.003 (0.0177) | 0.002 (0.0562) | Significance changed |
| FKBP4 | Chaperone | GSE113165 | 0.003 (0.0177) | 0.002 (0.0433) | No meaningful change |
| FKBP4 | Chaperone | GSE125892 | 0.003 (0.0177) | 0.003 (0.0527) | Significance changed |
| FKBP4 | Chaperone | GSE144304 | 0.003 (0.0177) | 0.002 (0.1119) | Significance changed |
| FKBP4 | Chaperone | GSE157712 | 0.003 (0.0177) | 0.003 (0.0174) | No meaningful change |
| FKBP4 | Chaperone | GSE159217 | 0.003 (0.0177) | 0.002 (0.0752) | Significance changed |
| FKBP4 | Chaperone | GSE167186 | 0.003 (0.0177) | 0.003 (0.0008) | No meaningful change |
| FKBP4 | Chaperone | GSE175495 | 0.003 (0.0177) | 0.003 (0.0047) | No meaningful change |
| FKBP4 | Chaperone | GSE181107 | 0.003 (0.0177) | 0.003 (0.0233) | No meaningful change |
| FKBP4 | Chaperone | GSE200398 | 0.003 (0.0177) | 0.003 (0.0127) | No meaningful change |
| FKBP4 | Chaperone | GSE208639 | 0.003 (0.0177) | 0.003 (0.0116) | No meaningful change |
| FKBP4 | Chaperone | GSE226646 | 0.003 (0.0177) | 0.003 (0.0078) | No meaningful change |
| FKBP4 | Chaperone | GSE242202 | 0.003 (0.0177) | 0.003 (0.0143) | No meaningful change |
| FKBP4 | Chaperone | GSE271606 | 0.003 (0.0177) | 0.003 (0.0381) | No meaningful change |
| FKBP4 | Chaperone | GSE287342 | 0.003 (0.0177) | 0.002 (0.0514) | Significance changed |
| FKBP5 | Chaperone | GSE111010 | 0.006 (0.0114) | 0.007 (0.0077) | No meaningful change |
| FKBP5 | Chaperone | GSE111016 | 0.006 (0.0114) | 0.006 (0.0123) | No meaningful change |
| FKBP5 | Chaperone | GSE113165 | 0.006 (0.0114) | 0.006 (0.0210) | No meaningful change |
| FKBP5 | Chaperone | GSE125892 | 0.006 (0.0114) | 0.006 (0.0146) | No meaningful change |
| FKBP5 | Chaperone | GSE144304 | 0.006 (0.0114) | 0.005 (0.0601) | Significance changed |
| FKBP5 | Chaperone | GSE157712 | 0.006 (0.0114) | 0.006 (0.0185) | No meaningful change |
| FKBP5 | Chaperone | GSE159217 | 0.006 (0.0114) | 0.008 (<0.0001) | No meaningful change |
| FKBP5 | Chaperone | GSE167186 | 0.006 (0.0114) | 0.006 (0.0578) | Significance changed |
| FKBP5 | Chaperone | GSE175495 | 0.006 (0.0114) | 0.006 (0.0284) | No meaningful change |
| FKBP5 | Chaperone | GSE181107 | 0.006 (0.0114) | 0.006 (0.0121) | No meaningful change |
| FKBP5 | Chaperone | GSE200398 | 0.006 (0.0114) | 0.006 (0.0160) | No meaningful change |
| FKBP5 | Chaperone | GSE208639 | 0.006 (0.0114) | 0.006 (0.0101) | No meaningful change |
| FKBP5 | Chaperone | GSE226646 | 0.006 (0.0114) | 0.006 (0.0116) | No meaningful change |
| FKBP5 | Chaperone | GSE242202 | 0.006 (0.0114) | 0.006 (0.0103) | No meaningful change |
| FKBP5 | Chaperone | GSE271606 | 0.006 (0.0114) | 0.006 (0.0108) | No meaningful change |
| FKBP5 | Chaperone | GSE287342 | 0.006 (0.0114) | 0.006 (0.0374) | No meaningful change |
| HSPA8 | Chaperone | GSE111010 | -0.004 (0.0015) | -0.004 (0.0026) | No meaningful change |
| HSPA8 | Chaperone | GSE111016 | -0.004 (0.0015) | -0.004 (0.0030) | No meaningful change |
| HSPA8 | Chaperone | GSE113165 | -0.004 (0.0015) | -0.003 (0.0188) | No meaningful change |
| HSPA8 | Chaperone | GSE125892 | -0.004 (0.0015) | -0.004 (0.0016) | No meaningful change |
| HSPA8 | Chaperone | GSE144304 | -0.004 (0.0015) | -0.004 (0.0018) | No meaningful change |
| HSPA8 | Chaperone | GSE157712 | -0.004 (0.0015) | -0.004 (0.0022) | No meaningful change |
| HSPA8 | Chaperone | GSE159217 | -0.004 (0.0015) | -0.003 (0.0079) | No meaningful change |
| HSPA8 | Chaperone | GSE167186 | -0.004 (0.0015) | -0.005 (0.0003) | No meaningful change |
| HSPA8 | Chaperone | GSE175495 | -0.004 (0.0015) | -0.004 (0.0063) | No meaningful change |
| HSPA8 | Chaperone | GSE181107 | -0.004 (0.0015) | -0.004 (0.0009) | No meaningful change |
| HSPA8 | Chaperone | GSE200398 | -0.004 (0.0015) | -0.004 (0.0025) | No meaningful change |
| HSPA8 | Chaperone | GSE208639 | -0.004 (0.0015) | -0.004 (0.0015) | No meaningful change |
| HSPA8 | Chaperone | GSE226646 | -0.004 (0.0015) | -0.004 (0.0018) | No meaningful change |
| HSPA8 | Chaperone | GSE242202 | -0.004 (0.0015) | -0.004 (0.0013) | No meaningful change |
| HSPA8 | Chaperone | GSE271606 | -0.004 (0.0015) | -0.004 (0.0014) | No meaningful change |
| HSPA8 | Chaperone | GSE287342 | -0.004 (0.0015) | -0.004 (0.0027) | No meaningful change |
| STIP1 | Chaperone | GSE111010 | -0.003 (0.2481) | -0.004 (0.1096) | No meaningful change |
| STIP1 | Chaperone | GSE111016 | -0.003 (0.2481) | -0.003 (0.3548) | No meaningful change |
| STIP1 | Chaperone | GSE113165 | -0.003 (0.2481) | -0.001 (0.5309) | No meaningful change |
| STIP1 | Chaperone | GSE125892 | -0.003 (0.2481) | -0.002 (0.3596) | No meaningful change |
| STIP1 | Chaperone | GSE144304 | -0.003 (0.2481) | -0.003 (0.2955) | No meaningful change |
| STIP1 | Chaperone | GSE157712 | -0.003 (0.2481) | -0.003 (0.2026) | No meaningful change |
| STIP1 | Chaperone | GSE159217 | -0.003 (0.2481) | -0.004 (0.2585) | No meaningful change |
| STIP1 | Chaperone | GSE167186 | -0.003 (0.2481) | -0.003 (0.2212) | No meaningful change |
| STIP1 | Chaperone | GSE175495 | -0.003 (0.2481) | -0.003 (0.4400) | No meaningful change |
| STIP1 | Chaperone | GSE181107 | -0.003 (0.2481) | -0.003 (0.2214) | No meaningful change |
| STIP1 | Chaperone | GSE200398 | -0.003 (0.2481) | -0.003 (0.3000) | No meaningful change |
| STIP1 | Chaperone | GSE208639 | -0.003 (0.2481) | -0.003 (0.2759) | No meaningful change |
| STIP1 | Chaperone | GSE226646 | -0.003 (0.2481) | -0.003 (0.2744) | No meaningful change |
| STIP1 | Chaperone | GSE242202 | -0.003 (0.2481) | -0.003 (0.2036) | No meaningful change |
| STIP1 | Chaperone | GSE271606 | -0.003 (0.2481) | -0.003 (0.2715) | No meaningful change |
| STIP1 | Chaperone | GSE287342 | -0.003 (0.2481) | -0.003 (0.2329) | No meaningful change |
| HSPB1 | Co-chaperone | GSE111010 | -0.003 (0.3805) | -0.003 (0.4250) | No meaningful change |
| HSPB1 | Co-chaperone | GSE111016 | -0.003 (0.3805) | -0.004 (0.2366) | No meaningful change |
| HSPB1 | Co-chaperone | GSE113165 | -0.003 (0.3805) | -0.003 (0.2863) | No meaningful change |
| HSPB1 | Co-chaperone | GSE125892 | -0.003 (0.3805) | -0.003 (0.2862) | No meaningful change |
| HSPB1 | Co-chaperone | GSE144304 | -0.003 (0.3805) | -0.003 (0.3277) | No meaningful change |
| HSPB1 | Co-chaperone | GSE157712 | -0.003 (0.3805) | -0.003 (0.4153) | No meaningful change |
| HSPB1 | Co-chaperone | GSE159217 | -0.003 (0.3805) | -0.003 (0.4122) | No meaningful change |
| HSPB1 | Co-chaperone | GSE167186 | -0.003 (0.3805) | -0.003 (0.4200) | No meaningful change |
| HSPB1 | Co-chaperone | GSE175495 | -0.003 (0.3805) | -0.002 (0.5333) | No meaningful change |
| HSPB1 | Co-chaperone | GSE181107 | -0.003 (0.3805) | -0.003 (0.3153) | No meaningful change |
| HSPB1 | Co-chaperone | GSE200398 | -0.003 (0.3805) | -0.003 (0.4084) | No meaningful change |
| HSPB1 | Co-chaperone | GSE208639 | -0.003 (0.3805) | -0.003 (0.4007) | No meaningful change |
| HSPB1 | Co-chaperone | GSE226646 | -0.003 (0.3805) | -0.001 (0.6727) | No meaningful change |
| HSPB1 | Co-chaperone | GSE242202 | -0.003 (0.3805) | -0.002 (0.5324) | No meaningful change |
| HSPB1 | Co-chaperone | GSE271606 | -0.003 (0.3805) | -0.001 (0.6591) | No meaningful change |
| HSPB1 | Co-chaperone | GSE287342 | -0.003 (0.3805) | -0.004 (0.2576) | No meaningful change |
| KPNA2 | Nuclear transport | GSE111010 | -0.008 (0.0072) | -0.009 (0.0028) | No meaningful change |
| KPNA2 | Nuclear transport | GSE111016 | -0.008 (0.0072) | -0.007 (0.0137) | No meaningful change |
| KPNA2 | Nuclear transport | GSE113165 | -0.008 (0.0072) | -0.005 (0.0096) | No meaningful change |
| KPNA2 | Nuclear transport | GSE125892 | -0.008 (0.0072) | -0.007 (0.0077) | No meaningful change |
| KPNA2 | Nuclear transport | GSE144304 | -0.008 (0.0072) | -0.008 (<0.0001) | No meaningful change |
| KPNA2 | Nuclear transport | GSE157712 | -0.008 (0.0072) | -0.007 (0.0124) | No meaningful change |
| KPNA2 | Nuclear transport | GSE159217 | -0.008 (0.0072) | -0.008 (0.0224) | No meaningful change |
| KPNA2 | Nuclear transport | GSE175495 | -0.008 (0.0072) | -0.008 (0.0184) | No meaningful change |
| KPNA2 | Nuclear transport | GSE181107 | -0.008 (0.0072) | -0.008 (0.0077) | No meaningful change |
| KPNA2 | Nuclear transport | GSE200398 | -0.008 (0.0072) | -0.008 (0.0138) | No meaningful change |
| KPNA2 | Nuclear transport | GSE208639 | -0.008 (0.0072) | -0.007 (0.0105) | No meaningful change |
| KPNA2 | Nuclear transport | GSE226646 | -0.008 (0.0072) | -0.008 (0.0067) | No meaningful change |
| KPNA2 | Nuclear transport | GSE242202 | -0.008 (0.0072) | -0.007 (0.0110) | No meaningful change |
| KPNA2 | Nuclear transport | GSE271606 | -0.008 (0.0072) | -0.007 (0.0101) | No meaningful change |
| KPNA2 | Nuclear transport | GSE287342 | -0.008 (0.0072) | -0.008 (<0.0001) | No meaningful change |
| KPNB1 | Nuclear transport | GSE111010 | -0.004 (<0.0001) | -0.004 (<0.0001) | No meaningful change |
| KPNB1 | Nuclear transport | GSE111016 | -0.004 (<0.0001) | -0.004 (0.0003) | No meaningful change |
| KPNB1 | Nuclear transport | GSE113165 | -0.004 (<0.0001) | -0.004 (<0.0001) | No meaningful change |
| KPNB1 | Nuclear transport | GSE125892 | -0.004 (<0.0001) | -0.004 (<0.0001) | No meaningful change |
| KPNB1 | Nuclear transport | GSE144304 | -0.004 (<0.0001) | -0.004 (0.0001) | No meaningful change |
| KPNB1 | Nuclear transport | GSE157712 | -0.004 (<0.0001) | -0.003 (<0.0001) | No meaningful change |
| KPNB1 | Nuclear transport | GSE159217 | -0.004 (<0.0001) | -0.004 (0.0004) | No meaningful change |
| KPNB1 | Nuclear transport | GSE167186 | -0.004 (<0.0001) | -0.004 (<0.0001) | No meaningful change |
| KPNB1 | Nuclear transport | GSE175495 | -0.004 (<0.0001) | -0.004 (0.0006) | No meaningful change |
| KPNB1 | Nuclear transport | GSE181107 | -0.004 (<0.0001) | -0.003 (<0.0001) | No meaningful change |
| KPNB1 | Nuclear transport | GSE200398 | -0.004 (<0.0001) | -0.004 (0.0002) | No meaningful change |
| KPNB1 | Nuclear transport | GSE208639 | -0.004 (<0.0001) | -0.003 (<0.0001) | No meaningful change |
| KPNB1 | Nuclear transport | GSE226646 | -0.004 (<0.0001) | -0.004 (0.0002) | No meaningful change |
| KPNB1 | Nuclear transport | GSE242202 | -0.004 (<0.0001) | -0.004 (<0.0001) | No meaningful change |
| KPNB1 | Nuclear transport | GSE271606 | -0.004 (<0.0001) | -0.004 (0.0001) | No meaningful change |
| KPNB1 | Nuclear transport | GSE287342 | -0.004 (<0.0001) | -0.004 (0.0002) | No meaningful change |
| RAN | Nuclear transport | GSE111010 | -0.001 (0.5920) | -0.001 (0.6824) | No meaningful change |
| RAN | Nuclear transport | GSE111016 | -0.001 (0.5920) | 0.000 (0.8384) | No meaningful change |
| RAN | Nuclear transport | GSE113165 | -0.001 (0.5920) | 0.000 (0.9928) | No meaningful change |
| RAN | Nuclear transport | GSE125892 | -0.001 (0.5920) | -0.001 (0.5485) | No meaningful change |
| RAN | Nuclear transport | GSE144304 | -0.001 (0.5920) | -0.001 (0.4325) | No meaningful change |
| RAN | Nuclear transport | GSE157712 | -0.001 (0.5920) | -0.001 (0.5922) | No meaningful change |
| RAN | Nuclear transport | GSE159217 | -0.001 (0.5920) | -0.001 (0.6336) | No meaningful change |
| RAN | Nuclear transport | GSE167186 | -0.001 (0.5920) | -0.001 (0.5189) | No meaningful change |
| RAN | Nuclear transport | GSE175495 | -0.001 (0.5920) | -0.001 (0.6588) | No meaningful change |
| RAN | Nuclear transport | GSE181107 | -0.001 (0.5920) | -0.001 (0.4147) | No meaningful change |
| RAN | Nuclear transport | GSE200398 | -0.001 (0.5920) | -0.001 (0.5410) | No meaningful change |
| RAN | Nuclear transport | GSE208639 | -0.001 (0.5920) | -0.001 (0.6331) | No meaningful change |
| RAN | Nuclear transport | GSE226646 | -0.001 (0.5920) | -0.001 (0.5514) | No meaningful change |
| RAN | Nuclear transport | GSE242202 | -0.001 (0.5920) | -0.001 (0.3458) | No meaningful change |
| RAN | Nuclear transport | GSE271606 | -0.001 (0.5920) | -0.001 (0.5586) | No meaningful change |
| RAN | Nuclear transport | GSE287342 | -0.001 (0.5920) | -0.001 (0.7803) | No meaningful change |
| KPNA1 | Nuclear transport | GSE111010 | 0.011 (0.0706) | 0.008 (0.1548) | No meaningful change |
| KPNA1 | Nuclear transport | GSE111016 | 0.011 (0.0706) | 0.012 (0.0700) | No meaningful change |
| KPNA1 | Nuclear transport | GSE125892 | 0.011 (0.0706) | 0.010 (0.1147) | No meaningful change |
| KPNA1 | Nuclear transport | GSE144304 | 0.011 (0.0706) | 0.010 (0.1468) | No meaningful change |
| KPNA1 | Nuclear transport | GSE159217 | 0.011 (0.0706) | 0.011 (0.1274) | No meaningful change |
| KPNA1 | Nuclear transport | GSE167186 | 0.011 (0.0706) | 0.015 (0.0352) | Significance changed |
| KPNA1 | Nuclear transport | GSE175495 | 0.011 (0.0706) | 0.015 (0.0224) | Significance changed |
| KPNA1 | Nuclear transport | GSE226646 | 0.011 (0.0706) | 0.013 (0.0581) | No meaningful change |
| KPNA1 | Nuclear transport | GSE242202 | 0.011 (0.0706) | 0.010 (0.0959) | No meaningful change |
| KPNA1 | Nuclear transport | GSE271606 | 0.011 (0.0706) | 0.009 (0.1142) | No meaningful change |
| KPNA5 | Nuclear transport | GSE111010 | -0.002 (0.0191) | -0.002 (0.0155) | No meaningful change |
| KPNA5 | Nuclear transport | GSE111016 | -0.002 (0.0191) | -0.002 (0.0414) | No meaningful change |
| KPNA5 | Nuclear transport | GSE113165 | -0.002 (0.0191) | -0.002 (0.0150) | No meaningful change |
| KPNA5 | Nuclear transport | GSE125892 | -0.002 (0.0191) | -0.002 (0.0188) | No meaningful change |
| KPNA5 | Nuclear transport | GSE144304 | -0.002 (0.0191) | -0.002 (0.0615) | Significance changed |
| KPNA5 | Nuclear transport | GSE157712 | -0.002 (0.0191) | -0.002 (0.0276) | No meaningful change |
| KPNA5 | Nuclear transport | GSE159217 | -0.002 (0.0191) | -0.002 (0.0164) | No meaningful change |
| KPNA5 | Nuclear transport | GSE167186 | -0.002 (0.0191) | -0.002 (0.0671) | Significance changed |
| KPNA5 | Nuclear transport | GSE175495 | -0.002 (0.0191) | -0.001 (0.0486) | No meaningful change |
| KPNA5 | Nuclear transport | GSE181107 | -0.002 (0.0191) | -0.002 (0.0189) | No meaningful change |
| KPNA5 | Nuclear transport | GSE200398 | -0.002 (0.0191) | -0.002 (0.0080) | No meaningful change |
| KPNA5 | Nuclear transport | GSE208639 | -0.002 (0.0191) | -0.002 (0.0153) | No meaningful change |
| KPNA5 | Nuclear transport | GSE226646 | -0.002 (0.0191) | -0.002 (0.0214) | No meaningful change |
| KPNA5 | Nuclear transport | GSE242202 | -0.002 (0.0191) | -0.002 (0.0214) | No meaningful change |
| KPNA5 | Nuclear transport | GSE271606 | -0.002 (0.0191) | -0.002 (0.0187) | No meaningful change |
| KPNA5 | Nuclear transport | GSE287342 | -0.002 (0.0191) | -0.002 (0.0211) | No meaningful change |
| KPNB2 | Nuclear transport | GSE111010 | 0.006 (0.0064) | 0.006 (0.0030) | No meaningful change |
| KPNB2 | Nuclear transport | GSE111016 | 0.006 (0.0064) | 0.006 (0.0119) | No meaningful change |
| KPNB2 | Nuclear transport | GSE113165 | 0.006 (0.0064) | 0.006 (0.0092) | No meaningful change |
| KPNB2 | Nuclear transport | GSE144304 | 0.006 (0.0064) | 0.004 (0.0486) | No meaningful change |
| KPNB2 | Nuclear transport | GSE157712 | 0.006 (0.0064) | 0.005 (0.0258) | No meaningful change |
| KPNB2 | Nuclear transport | GSE159217 | 0.006 (0.0064) | 0.007 (0.0021) | No meaningful change |
| KPNB2 | Nuclear transport | GSE181107 | 0.006 (0.0064) | 0.005 (0.0101) | No meaningful change |
| KPNB2 | Nuclear transport | GSE200398 | 0.006 (0.0064) | 0.005 (0.0138) | No meaningful change |
| KPNB2 | Nuclear transport | GSE208639 | 0.006 (0.0064) | 0.006 (0.0011) | No meaningful change |
| KPNB2 | Nuclear transport | GSE242202 | 0.006 (0.0064) | 0.006 (0.0088) | No meaningful change |
| KPNB2 | Nuclear transport | GSE271606 | 0.006 (0.0064) | 0.005 (0.0121) | No meaningful change |
| NCOA2 | Coactivator | GSE111010 | 0.002 (0.1076) | 0.002 (0.1191) | No meaningful change |
| NCOA2 | Coactivator | GSE111016 | 0.002 (0.1076) | 0.002 (0.1258) | No meaningful change |
| NCOA2 | Coactivator | GSE113165 | 0.002 (0.1076) | 0.001 (0.1930) | No meaningful change |
| NCOA2 | Coactivator | GSE125892 | 0.002 (0.1076) | 0.002 (0.0988) | No meaningful change |
| NCOA2 | Coactivator | GSE144304 | 0.002 (0.1076) | 0.001 (0.3138) | No meaningful change |
| NCOA2 | Coactivator | GSE157712 | 0.002 (0.1076) | 0.002 (0.1641) | No meaningful change |
| NCOA2 | Coactivator | GSE159217 | 0.002 (0.1076) | 0.002 (0.1277) | No meaningful change |
| NCOA2 | Coactivator | GSE167186 | 0.002 (0.1076) | 0.002 (0.1297) | No meaningful change |
| NCOA2 | Coactivator | GSE175495 | 0.002 (0.1076) | 0.002 (0.0098) | Significance changed |
| NCOA2 | Coactivator | GSE181107 | 0.002 (0.1076) | 0.002 (0.0849) | No meaningful change |
| NCOA2 | Coactivator | GSE200398 | 0.002 (0.1076) | 0.001 (0.2429) | No meaningful change |
| NCOA2 | Coactivator | GSE208639 | 0.002 (0.1076) | 0.002 (0.1069) | No meaningful change |
| NCOA2 | Coactivator | GSE226646 | 0.002 (0.1076) | 0.002 (0.1050) | No meaningful change |
| NCOA2 | Coactivator | GSE242202 | 0.002 (0.1076) | 0.002 (0.1206) | No meaningful change |
| NCOA2 | Coactivator | GSE271606 | 0.002 (0.1076) | 0.002 (0.1039) | No meaningful change |
| NCOA2 | Coactivator | GSE287342 | 0.002 (0.1076) | 0.002 (0.0626) | No meaningful change |
| NCOA3 | Coactivator | GSE111010 | -0.001 (0.3691) | -0.001 (0.4849) | No meaningful change |
| NCOA3 | Coactivator | GSE111016 | -0.001 (0.3691) | -0.001 (0.3082) | No meaningful change |
| NCOA3 | Coactivator | GSE113165 | -0.001 (0.3691) | -0.001 (0.4241) | No meaningful change |
| NCOA3 | Coactivator | GSE125892 | -0.001 (0.3691) | -0.001 (0.3944) | No meaningful change |
| NCOA3 | Coactivator | GSE144304 | -0.001 (0.3691) | -0.001 (0.4046) | No meaningful change |
| NCOA3 | Coactivator | GSE157712 | -0.001 (0.3691) | -0.001 (0.4118) | No meaningful change |
| NCOA3 | Coactivator | GSE159217 | -0.001 (0.3691) | -0.001 (0.2296) | No meaningful change |
| NCOA3 | Coactivator | GSE167186 | -0.001 (0.3691) | -0.001 (0.6545) | No meaningful change |
| NCOA3 | Coactivator | GSE175495 | -0.001 (0.3691) | -0.001 (0.6165) | No meaningful change |
| NCOA3 | Coactivator | GSE181107 | -0.001 (0.3691) | -0.001 (0.3673) | No meaningful change |
| NCOA3 | Coactivator | GSE200398 | -0.001 (0.3691) | -0.001 (0.4682) | No meaningful change |
| NCOA3 | Coactivator | GSE208639 | -0.001 (0.3691) | -0.001 (0.3812) | No meaningful change |
| NCOA3 | Coactivator | GSE226646 | -0.001 (0.3691) | -0.001 (0.3914) | No meaningful change |
| NCOA3 | Coactivator | GSE242202 | -0.001 (0.3691) | -0.001 (0.3979) | No meaningful change |
| NCOA3 | Coactivator | GSE271606 | -0.001 (0.3691) | -0.001 (0.4054) | No meaningful change |
| NCOA3 | Coactivator | GSE287342 | -0.001 (0.3691) | -0.002 (0.0687) | No meaningful change |
| CREBBP | Coactivator | GSE111010 | 0.004 (0.0009) | 0.004 (0.0015) | No meaningful change |
| CREBBP | Coactivator | GSE111016 | 0.004 (0.0009) | 0.003 (0.0023) | No meaningful change |
| CREBBP | Coactivator | GSE113165 | 0.004 (0.0009) | 0.003 (0.0030) | No meaningful change |
| CREBBP | Coactivator | GSE125892 | 0.004 (0.0009) | 0.004 (0.0010) | No meaningful change |
| CREBBP | Coactivator | GSE144304 | 0.004 (0.0009) | 0.003 (0.0033) | No meaningful change |
| CREBBP | Coactivator | GSE157712 | 0.004 (0.0009) | 0.004 (0.0011) | No meaningful change |
| CREBBP | Coactivator | GSE159217 | 0.004 (0.0009) | 0.004 (0.0020) | No meaningful change |
| CREBBP | Coactivator | GSE167186 | 0.004 (0.0009) | 0.004 (<0.0001) | No meaningful change |
| CREBBP | Coactivator | GSE175495 | 0.004 (0.0009) | 0.004 (0.0023) | No meaningful change |
| CREBBP | Coactivator | GSE181107 | 0.004 (0.0009) | 0.004 (0.0011) | No meaningful change |
| CREBBP | Coactivator | GSE200398 | 0.004 (0.0009) | 0.004 (0.0007) | No meaningful change |
| CREBBP | Coactivator | GSE208639 | 0.004 (0.0009) | 0.004 (0.0027) | No meaningful change |
| CREBBP | Coactivator | GSE226646 | 0.004 (0.0009) | 0.004 (0.0013) | No meaningful change |
| CREBBP | Coactivator | GSE242202 | 0.004 (0.0009) | 0.004 (0.0003) | No meaningful change |
| CREBBP | Coactivator | GSE271606 | 0.004 (0.0009) | 0.004 (0.0008) | No meaningful change |
| CREBBP | Coactivator | GSE287342 | 0.004 (0.0009) | 0.004 (0.0015) | No meaningful change |
| EP300 | Coactivator | GSE111010 | 0.002 (0.0007) | 0.002 (0.0012) | No meaningful change |
| EP300 | Coactivator | GSE111016 | 0.002 (0.0007) | 0.002 (0.0018) | No meaningful change |
| EP300 | Coactivator | GSE113165 | 0.002 (0.0007) | 0.002 (0.0007) | No meaningful change |
| EP300 | Coactivator | GSE125892 | 0.002 (0.0007) | 0.002 (0.0007) | No meaningful change |
| EP300 | Coactivator | GSE144304 | 0.002 (0.0007) | 0.002 (0.0061) | No meaningful change |
| EP300 | Coactivator | GSE157712 | 0.002 (0.0007) | 0.002 (0.0007) | No meaningful change |
| EP300 | Coactivator | GSE159217 | 0.002 (0.0007) | 0.002 (0.0024) | No meaningful change |
| EP300 | Coactivator | GSE167186 | 0.002 (0.0007) | 0.003 (0.0011) | No meaningful change |
| EP300 | Coactivator | GSE175495 | 0.002 (0.0007) | 0.002 (0.0008) | No meaningful change |
| EP300 | Coactivator | GSE181107 | 0.002 (0.0007) | 0.002 (0.0008) | No meaningful change |
| EP300 | Coactivator | GSE200398 | 0.002 (0.0007) | 0.002 (0.0007) | No meaningful change |
| EP300 | Coactivator | GSE208639 | 0.002 (0.0007) | 0.002 (0.0006) | No meaningful change |
| EP300 | Coactivator | GSE226646 | 0.002 (0.0007) | 0.002 (0.0007) | No meaningful change |
| EP300 | Coactivator | GSE242202 | 0.002 (0.0007) | 0.002 (0.0006) | No meaningful change |
| EP300 | Coactivator | GSE271606 | 0.002 (0.0007) | 0.002 (0.0007) | No meaningful change |
| EP300 | Coactivator | GSE287342 | 0.002 (0.0007) | 0.002 (0.0016) | No meaningful change |
| NCOR1 | Corepressor | GSE111010 | 0.001 (0.0048) | 0.001 (0.0072) | No meaningful change |
| NCOR1 | Corepressor | GSE111016 | 0.001 (0.0048) | 0.001 (0.0088) | No meaningful change |
| NCOR1 | Corepressor | GSE113165 | 0.001 (0.0048) | 0.001 (0.0188) | No meaningful change |
| NCOR1 | Corepressor | GSE125892 | 0.001 (0.0048) | 0.001 (0.0048) | No meaningful change |
| NCOR1 | Corepressor | GSE144304 | 0.001 (0.0048) | 0.001 (0.0052) | No meaningful change |
| NCOR1 | Corepressor | GSE157712 | 0.001 (0.0048) | 0.001 (0.0044) | No meaningful change |
| NCOR1 | Corepressor | GSE159217 | 0.001 (0.0048) | 0.002 (0.0137) | No meaningful change |
| NCOR1 | Corepressor | GSE167186 | 0.001 (0.0048) | 0.002 (0.0012) | No meaningful change |
| NCOR1 | Corepressor | GSE175495 | 0.001 (0.0048) | 0.002 (0.0160) | No meaningful change |
| NCOR1 | Corepressor | GSE181107 | 0.001 (0.0048) | 0.001 (0.0048) | No meaningful change |
| NCOR1 | Corepressor | GSE200398 | 0.001 (0.0048) | 0.001 (0.0045) | No meaningful change |
| NCOR1 | Corepressor | GSE208639 | 0.001 (0.0048) | 0.001 (0.0045) | No meaningful change |
| NCOR1 | Corepressor | GSE226646 | 0.001 (0.0048) | 0.001 (0.0049) | No meaningful change |
| NCOR1 | Corepressor | GSE242202 | 0.001 (0.0048) | 0.001 (0.0055) | No meaningful change |
| NCOR1 | Corepressor | GSE271606 | 0.001 (0.0048) | 0.001 (0.0045) | No meaningful change |
| NCOR1 | Corepressor | GSE287342 | 0.001 (0.0048) | 0.001 (0.0046) | No meaningful change |
| NCOR2 | Corepressor | GSE111010 | 0.001 (0.2817) | 0.001 (0.3003) | No meaningful change |
| NCOR2 | Corepressor | GSE111016 | 0.001 (0.2817) | 0.001 (0.3094) | No meaningful change |
| NCOR2 | Corepressor | GSE113165 | 0.001 (0.2817) | 0.001 (0.3039) | No meaningful change |
| NCOR2 | Corepressor | GSE125892 | 0.001 (0.2817) | 0.001 (0.2967) | No meaningful change |
| NCOR2 | Corepressor | GSE144304 | 0.001 (0.2817) | 0.001 (0.3819) | No meaningful change |
| NCOR2 | Corepressor | GSE157712 | 0.001 (0.2817) | 0.001 (0.1764) | No meaningful change |
| NCOR2 | Corepressor | GSE159217 | 0.001 (0.2817) | 0.001 (0.1948) | No meaningful change |
| NCOR2 | Corepressor | GSE167186 | 0.001 (0.2817) | 0.001 (0.5645) | No meaningful change |
| NCOR2 | Corepressor | GSE175495 | 0.001 (0.2817) | 0.001 (0.2472) | No meaningful change |
| NCOR2 | Corepressor | GSE181107 | 0.001 (0.2817) | 0.001 (0.2740) | No meaningful change |
| NCOR2 | Corepressor | GSE200398 | 0.001 (0.2817) | 0.001 (0.3455) | No meaningful change |
| NCOR2 | Corepressor | GSE208639 | 0.001 (0.2817) | 0.001 (0.3735) | No meaningful change |
| NCOR2 | Corepressor | GSE226646 | 0.001 (0.2817) | 0.001 (0.3084) | No meaningful change |
| NCOR2 | Corepressor | GSE242202 | 0.001 (0.2817) | 0.001 (0.2976) | No meaningful change |
| NCOR2 | Corepressor | GSE271606 | 0.001 (0.2817) | 0.001 (0.2856) | No meaningful change |
| NCOR2 | Corepressor | GSE287342 | 0.001 (0.2817) | 0.001 (0.3152) | No meaningful change |
| GATA2 | Pioneer factor | GSE111010 | 0.000 (0.8634) | -0.001 (0.6207) | No meaningful change |
| GATA2 | Pioneer factor | GSE111016 | 0.000 (0.8634) | 0.000 (0.9805) | No meaningful change |
| GATA2 | Pioneer factor | GSE113165 | 0.000 (0.8634) | 0.000 (0.8858) | No meaningful change |
| GATA2 | Pioneer factor | GSE144304 | 0.000 (0.8634) | 0.000 (0.8780) | No meaningful change |
| GATA2 | Pioneer factor | GSE157712 | 0.000 (0.8634) | -0.003 (0.2233) | No meaningful change |
| GATA2 | Pioneer factor | GSE159217 | 0.000 (0.8634) | 0.001 (0.7514) | No meaningful change |
| GATA2 | Pioneer factor | GSE175495 | 0.000 (0.8634) | -0.001 (0.8028) | No meaningful change |
| GATA2 | Pioneer factor | GSE181107 | 0.000 (0.8634) | 0.000 (0.8870) | No meaningful change |
| GATA2 | Pioneer factor | GSE200398 | 0.000 (0.8634) | 0.000 (0.9305) | No meaningful change |
| GATA2 | Pioneer factor | GSE208639 | 0.000 (0.8634) | 0.000 (0.9258) | No meaningful change |
| GATA2 | Pioneer factor | GSE242202 | 0.000 (0.8634) | 0.000 (0.9153) | No meaningful change |
| GATA2 | Pioneer factor | GSE271606 | 0.000 (0.8634) | 0.000 (0.8815) | No meaningful change |
| GATA2 | Pioneer factor | GSE287342 | 0.000 (0.8634) | -0.001 (0.6743) | No meaningful change |
| MED1 | Mediator complex | GSE111010 | 0.001 (0.4685) | 0.001 (0.4767) | No meaningful change |
| MED1 | Mediator complex | GSE111016 | 0.001 (0.4685) | 0.001 (0.4698) | No meaningful change |
| MED1 | Mediator complex | GSE113165 | 0.001 (0.4685) | 0.001 (0.6155) | No meaningful change |
| MED1 | Mediator complex | GSE125892 | 0.001 (0.4685) | 0.001 (0.4223) | No meaningful change |
| MED1 | Mediator complex | GSE167186 | 0.001 (0.4685) | 0.000 (0.7562) | No meaningful change |
| MED1 | Mediator complex | GSE175495 | 0.001 (0.4685) | 0.002 (0.0401) | Significance changed |
| MED1 | Mediator complex | GSE226646 | 0.001 (0.4685) | 0.001 (0.4677) | No meaningful change |
| MED1 | Mediator complex | GSE242202 | 0.001 (0.4685) | 0.001 (0.4438) | No meaningful change |
| MED1 | Mediator complex | GSE271606 | 0.001 (0.4685) | 0.001 (0.4742) | No meaningful change |
| MED1 | Mediator complex | GSE287342 | 0.001 (0.4685) | 0.001 (0.4155) | No meaningful change |
| SMARCA4 | Chromatin remodeler | GSE111010 | -0.002 (0.0210) | -0.002 (0.0076) | No meaningful change |
| SMARCA4 | Chromatin remodeler | GSE111016 | -0.002 (0.0210) | -0.002 (0.0997) | Significance changed |
| SMARCA4 | Chromatin remodeler | GSE113165 | -0.002 (0.0210) | -0.002 (0.0162) | No meaningful change |
| SMARCA4 | Chromatin remodeler | GSE125892 | -0.002 (0.0210) | -0.002 (0.0098) | No meaningful change |
| SMARCA4 | Chromatin remodeler | GSE144304 | -0.002 (0.0210) | -0.001 (0.2166) | Significance changed |
| SMARCA4 | Chromatin remodeler | GSE157712 | -0.002 (0.0210) | -0.002 (0.0822) | Significance changed |
| SMARCA4 | Chromatin remodeler | GSE159217 | -0.002 (0.0210) | -0.002 (0.0279) | No meaningful change |
| SMARCA4 | Chromatin remodeler | GSE167186 | -0.002 (0.0210) | -0.002 (0.0323) | No meaningful change |
| SMARCA4 | Chromatin remodeler | GSE175495 | -0.002 (0.0210) | -0.002 (0.0128) | No meaningful change |
| SMARCA4 | Chromatin remodeler | GSE181107 | -0.002 (0.0210) | -0.002 (0.0061) | No meaningful change |
| SMARCA4 | Chromatin remodeler | GSE200398 | -0.002 (0.0210) | -0.002 (0.0113) | No meaningful change |
| SMARCA4 | Chromatin remodeler | GSE208639 | -0.002 (0.0210) | -0.002 (0.0238) | No meaningful change |
| SMARCA4 | Chromatin remodeler | GSE226646 | -0.002 (0.0210) | -0.002 (0.0074) | No meaningful change |
| SMARCA4 | Chromatin remodeler | GSE242202 | -0.002 (0.0210) | -0.002 (0.0594) | Significance changed |
| SMARCA4 | Chromatin remodeler | GSE271606 | -0.002 (0.0210) | -0.002 (0.0185) | No meaningful change |
| SMARCA4 | Chromatin remodeler | GSE287342 | -0.002 (0.0210) | -0.001 (0.1750) | Significance changed |
| KDM1A | Chromatin modifier | GSE111010 | 0.000 (0.9836) | 0.000 (0.8774) | No meaningful change |
| KDM1A | Chromatin modifier | GSE111016 | 0.000 (0.9836) | 0.000 (0.8744) | No meaningful change |
| KDM1A | Chromatin modifier | GSE113165 | 0.000 (0.9836) | 0.000 (0.7870) | No meaningful change |
| KDM1A | Chromatin modifier | GSE125892 | 0.000 (0.9836) | 0.000 (0.9928) | No meaningful change |
| KDM1A | Chromatin modifier | GSE144304 | 0.000 (0.9836) | -0.001 (0.0259) | Significance changed |
| KDM1A | Chromatin modifier | GSE157712 | 0.000 (0.9836) | 0.000 (0.9125) | No meaningful change |
| KDM1A | Chromatin modifier | GSE159217 | 0.000 (0.9836) | 0.001 (0.6427) | No meaningful change |
| KDM1A | Chromatin modifier | GSE167186 | 0.000 (0.9836) | 0.000 (0.6757) | No meaningful change |
| KDM1A | Chromatin modifier | GSE175495 | 0.000 (0.9836) | 0.001 (0.5550) | No meaningful change |
| KDM1A | Chromatin modifier | GSE181107 | 0.000 (0.9836) | 0.000 (0.8843) | No meaningful change |
| KDM1A | Chromatin modifier | GSE200398 | 0.000 (0.9836) | 0.000 (0.9728) | No meaningful change |
| KDM1A | Chromatin modifier | GSE208639 | 0.000 (0.9836) | 0.000 (0.9788) | No meaningful change |
| KDM1A | Chromatin modifier | GSE226646 | 0.000 (0.9836) | 0.000 (0.9682) | No meaningful change |
| KDM1A | Chromatin modifier | GSE242202 | 0.000 (0.9836) | 0.000 (0.9835) | No meaningful change |
| KDM1A | Chromatin modifier | GSE271606 | 0.000 (0.9836) | 0.000 (0.8400) | No meaningful change |
| KDM1A | Chromatin modifier | GSE287342 | 0.000 (0.9836) | 0.000 (0.8497) | No meaningful change |
| STUB1 | Ubiquitination | GSE111010 | -0.002 (0.0203) | -0.002 (0.0174) | No meaningful change |
| STUB1 | Ubiquitination | GSE111016 | -0.002 (0.0203) | -0.002 (0.0210) | No meaningful change |
| STUB1 | Ubiquitination | GSE125892 | -0.002 (0.0203) | -0.002 (0.0246) | No meaningful change |
| STUB1 | Ubiquitination | GSE144304 | -0.002 (0.0203) | -0.002 (0.0229) | No meaningful change |
| STUB1 | Ubiquitination | GSE157712 | -0.002 (0.0203) | -0.002 (0.0217) | No meaningful change |
| STUB1 | Ubiquitination | GSE159217 | -0.002 (0.0203) | -0.002 (0.0743) | Significance changed |
| STUB1 | Ubiquitination | GSE167186 | -0.002 (0.0203) | -0.003 (0.0093) | No meaningful change |
| STUB1 | Ubiquitination | GSE175495 | -0.002 (0.0203) | -0.003 (0.0528) | Significance changed |
| STUB1 | Ubiquitination | GSE181107 | -0.002 (0.0203) | -0.002 (0.0261) | No meaningful change |
| STUB1 | Ubiquitination | GSE200398 | -0.002 (0.0203) | -0.002 (0.0367) | No meaningful change |
| STUB1 | Ubiquitination | GSE208639 | -0.002 (0.0203) | -0.002 (0.0350) | No meaningful change |
| STUB1 | Ubiquitination | GSE226646 | -0.002 (0.0203) | -0.002 (0.0195) | No meaningful change |
| STUB1 | Ubiquitination | GSE242202 | -0.002 (0.0203) | -0.002 (0.0243) | No meaningful change |
| STUB1 | Ubiquitination | GSE271606 | -0.002 (0.0203) | -0.002 (0.0243) | No meaningful change |
| UBC | Ubiquitination | GSE111010 | -0.002 (0.1572) | -0.002 (0.1801) | No meaningful change |
| UBC | Ubiquitination | GSE111016 | -0.002 (0.1572) | -0.002 (0.1306) | No meaningful change |
| UBC | Ubiquitination | GSE113165 | -0.002 (0.1572) | -0.001 (0.4310) | No meaningful change |
| UBC | Ubiquitination | GSE144304 | -0.002 (0.1572) | -0.002 (0.2635) | No meaningful change |
| UBC | Ubiquitination | GSE157712 | -0.002 (0.1572) | -0.003 (0.0677) | No meaningful change |
| UBC | Ubiquitination | GSE159217 | -0.002 (0.1572) | -0.003 (0.1482) | No meaningful change |
| UBC | Ubiquitination | GSE167186 | -0.002 (0.1572) | -0.002 (0.1892) | No meaningful change |
| UBC | Ubiquitination | GSE175495 | -0.002 (0.1572) | -0.002 (0.2393) | No meaningful change |
| UBC | Ubiquitination | GSE181107 | -0.002 (0.1572) | -0.002 (0.1819) | No meaningful change |
| UBC | Ubiquitination | GSE200398 | -0.002 (0.1572) | -0.002 (0.1768) | No meaningful change |
| UBC | Ubiquitination | GSE208639 | -0.002 (0.1572) | -0.001 (0.3927) | No meaningful change |
| UBC | Ubiquitination | GSE226646 | -0.002 (0.1572) | -0.003 (0.0886) | No meaningful change |
| UBC | Ubiquitination | GSE242202 | -0.002 (0.1572) | -0.003 (0.1652) | No meaningful change |
| UBC | Ubiquitination | GSE271606 | -0.002 (0.1572) | -0.003 (0.2263) | No meaningful change |
| UBC | Ubiquitination | GSE287342 | -0.002 (0.1572) | -0.003 (0.1994) | No meaningful change |
| UBE2I | SUMOylation | GSE111010 | 0.002 (0.2086) | 0.002 (0.1613) | No meaningful change |
| UBE2I | SUMOylation | GSE111016 | 0.002 (0.2086) | 0.001 (0.3313) | No meaningful change |
| UBE2I | SUMOylation | GSE113165 | 0.002 (0.2086) | 0.002 (0.1590) | No meaningful change |
| UBE2I | SUMOylation | GSE125892 | 0.002 (0.2086) | 0.001 (0.2434) | No meaningful change |
| UBE2I | SUMOylation | GSE144304 | 0.002 (0.2086) | 0.001 (0.2461) | No meaningful change |
| UBE2I | SUMOylation | GSE157712 | 0.002 (0.2086) | 0.003 (0.0390) | Significance changed |
| UBE2I | SUMOylation | GSE159217 | 0.002 (0.2086) | 0.002 (0.2387) | No meaningful change |
| UBE2I | SUMOylation | GSE167186 | 0.002 (0.2086) | 0.002 (0.2791) | No meaningful change |
| UBE2I | SUMOylation | GSE175495 | 0.002 (0.2086) | 0.002 (0.2519) | No meaningful change |
| UBE2I | SUMOylation | GSE181107 | 0.002 (0.2086) | 0.001 (0.2811) | No meaningful change |
| UBE2I | SUMOylation | GSE200398 | 0.002 (0.2086) | 0.002 (0.2467) | No meaningful change |
| UBE2I | SUMOylation | GSE208639 | 0.002 (0.2086) | 0.002 (0.1955) | No meaningful change |
| UBE2I | SUMOylation | GSE226646 | 0.002 (0.2086) | 0.002 (0.1034) | No meaningful change |
| UBE2I | SUMOylation | GSE242202 | 0.002 (0.2086) | 0.002 (0.1900) | No meaningful change |
| UBE2I | SUMOylation | GSE271606 | 0.002 (0.2086) | 0.002 (0.2193) | No meaningful change |
| UBE2I | SUMOylation | GSE287342 | 0.002 (0.2086) | 0.002 (0.3446) | No meaningful change |
| SPOP | Ubiquitination | GSE111010 | 0.004 (0.0112) | 0.004 (0.0119) | No meaningful change |
| SPOP | Ubiquitination | GSE111016 | 0.004 (0.0112) | 0.004 (0.0139) | No meaningful change |
| SPOP | Ubiquitination | GSE113165 | 0.004 (0.0112) | 0.003 (0.0375) | No meaningful change |
| SPOP | Ubiquitination | GSE125892 | 0.004 (0.0112) | 0.004 (0.0131) | No meaningful change |
| SPOP | Ubiquitination | GSE144304 | 0.004 (0.0112) | 0.003 (0.0258) | No meaningful change |
| SPOP | Ubiquitination | GSE157712 | 0.004 (0.0112) | 0.004 (0.0085) | No meaningful change |
| SPOP | Ubiquitination | GSE159217 | 0.004 (0.0112) | 0.005 (0.0063) | No meaningful change |
| SPOP | Ubiquitination | GSE167186 | 0.004 (0.0112) | 0.005 (0.0005) | No meaningful change |
| SPOP | Ubiquitination | GSE175495 | 0.004 (0.0112) | 0.004 (0.0278) | No meaningful change |
| SPOP | Ubiquitination | GSE181107 | 0.004 (0.0112) | 0.004 (0.0067) | No meaningful change |
| SPOP | Ubiquitination | GSE200398 | 0.004 (0.0112) | 0.004 (0.0150) | No meaningful change |
| SPOP | Ubiquitination | GSE208639 | 0.004 (0.0112) | 0.004 (0.0115) | No meaningful change |
| SPOP | Ubiquitination | GSE226646 | 0.004 (0.0112) | 0.004 (0.0117) | No meaningful change |
| SPOP | Ubiquitination | GSE242202 | 0.004 (0.0112) | 0.004 (0.0105) | No meaningful change |
| SPOP | Ubiquitination | GSE271606 | 0.004 (0.0112) | 0.004 (0.0073) | No meaningful change |
| SPOP | Ubiquitination | GSE287342 | 0.004 (0.0112) | 0.003 (0.0316) | No meaningful change |
| MDM2 | Ubiquitination | GSE111010 | 0.002 (0.0049) | 0.002 (0.0071) | No meaningful change |
| MDM2 | Ubiquitination | GSE111016 | 0.002 (0.0049) | 0.002 (0.0054) | No meaningful change |
| MDM2 | Ubiquitination | GSE113165 | 0.002 (0.0049) | 0.002 (0.0250) | No meaningful change |
| MDM2 | Ubiquitination | GSE125892 | 0.002 (0.0049) | 0.002 (0.0050) | No meaningful change |
| MDM2 | Ubiquitination | GSE144304 | 0.002 (0.0049) | 0.002 (0.0590) | Significance changed |
| MDM2 | Ubiquitination | GSE157712 | 0.002 (0.0049) | 0.002 (0.0058) | No meaningful change |
| MDM2 | Ubiquitination | GSE159217 | 0.002 (0.0049) | 0.002 (0.0903) | Significance changed |
| MDM2 | Ubiquitination | GSE167186 | 0.002 (0.0049) | 0.002 (0.0583) | Significance changed |
| MDM2 | Ubiquitination | GSE175495 | 0.002 (0.0049) | 0.003 (<0.0001) | No meaningful change |
| MDM2 | Ubiquitination | GSE181107 | 0.002 (0.0049) | 0.002 (0.0050) | No meaningful change |
| MDM2 | Ubiquitination | GSE200398 | 0.002 (0.0049) | 0.003 (0.0014) | No meaningful change |
| MDM2 | Ubiquitination | GSE208639 | 0.002 (0.0049) | 0.003 (0.0043) | No meaningful change |
| MDM2 | Ubiquitination | GSE226646 | 0.002 (0.0049) | 0.002 (0.0059) | No meaningful change |
| MDM2 | Ubiquitination | GSE242202 | 0.002 (0.0049) | 0.003 (0.0047) | No meaningful change |
| MDM2 | Ubiquitination | GSE271606 | 0.002 (0.0049) | 0.002 (0.0053) | No meaningful change |
| MDM2 | Ubiquitination | GSE287342 | 0.002 (0.0049) | 0.003 (0.0036) | No meaningful change |
| AKT1 | Kinase | GSE111010 | -0.002 (0.2121) | -0.002 (0.2906) | No meaningful change |
| AKT1 | Kinase | GSE111016 | -0.002 (0.2121) | -0.002 (0.3120) | No meaningful change |
| AKT1 | Kinase | GSE113165 | -0.002 (0.2121) | -0.001 (0.0292) | Significance changed |
| AKT1 | Kinase | GSE125892 | -0.002 (0.2121) | -0.002 (0.2224) | No meaningful change |
| AKT1 | Kinase | GSE144304 | -0.002 (0.2121) | -0.002 (0.2185) | No meaningful change |
| AKT1 | Kinase | GSE157712 | -0.002 (0.2121) | -0.003 (0.0781) | No meaningful change |
| AKT1 | Kinase | GSE159217 | -0.002 (0.2121) | -0.002 (0.2484) | No meaningful change |
| AKT1 | Kinase | GSE175495 | -0.002 (0.2121) | -0.002 (0.2841) | No meaningful change |
| AKT1 | Kinase | GSE181107 | -0.002 (0.2121) | -0.002 (0.1985) | No meaningful change |
| AKT1 | Kinase | GSE200398 | -0.002 (0.2121) | -0.003 (0.1090) | No meaningful change |
| AKT1 | Kinase | GSE208639 | -0.002 (0.2121) | -0.002 (0.0410) | Significance changed |
| AKT1 | Kinase | GSE226646 | -0.002 (0.2121) | -0.002 (0.0769) | No meaningful change |
| AKT1 | Kinase | GSE242202 | -0.002 (0.2121) | -0.002 (0.1513) | No meaningful change |
| AKT1 | Kinase | GSE271606 | -0.002 (0.2121) | -0.002 (0.1954) | No meaningful change |
| AKT1 | Kinase | GSE287342 | -0.002 (0.2121) | -0.001 (0.0788) | No meaningful change |
| MAPK1 | Kinase | GSE111010 | -0.001 (0.1439) | -0.001 (0.1544) | No meaningful change |
| MAPK1 | Kinase | GSE111016 | -0.001 (0.1439) | -0.001 (0.1518) | No meaningful change |
| MAPK1 | Kinase | GSE113165 | -0.001 (0.1439) | 0.000 (0.4484) | No meaningful change |
| MAPK1 | Kinase | GSE125892 | -0.001 (0.1439) | -0.001 (0.1792) | No meaningful change |
| MAPK1 | Kinase | GSE144304 | -0.001 (0.1439) | -0.001 (0.1731) | No meaningful change |
| MAPK1 | Kinase | GSE157712 | -0.001 (0.1439) | -0.001 (0.1717) | No meaningful change |
| MAPK1 | Kinase | GSE159217 | -0.001 (0.1439) | -0.001 (0.0456) | Significance changed |
| MAPK1 | Kinase | GSE167186 | -0.001 (0.1439) | -0.001 (0.1062) | No meaningful change |
| MAPK1 | Kinase | GSE175495 | -0.001 (0.1439) | -0.001 (0.2913) | No meaningful change |
| MAPK1 | Kinase | GSE181107 | -0.001 (0.1439) | -0.001 (0.1133) | No meaningful change |
| MAPK1 | Kinase | GSE200398 | -0.001 (0.1439) | -0.001 (0.1441) | No meaningful change |
| MAPK1 | Kinase | GSE208639 | -0.001 (0.1439) | -0.001 (0.1777) | No meaningful change |
| MAPK1 | Kinase | GSE226646 | -0.001 (0.1439) | -0.001 (0.1360) | No meaningful change |
| MAPK1 | Kinase | GSE242202 | -0.001 (0.1439) | -0.001 (0.1256) | No meaningful change |
| MAPK1 | Kinase | GSE271606 | -0.001 (0.1439) | -0.001 (0.1359) | No meaningful change |
| MAPK1 | Kinase | GSE287342 | -0.001 (0.1439) | -0.001 (0.1463) | No meaningful change |
| MYOD1 | Muscle differentiation | GSE125892 | 0.003 (0.2905) | 0.003 (0.2889) | No meaningful change |
| MYOD1 | Muscle differentiation | GSE167186 | 0.003 (0.2905) | -0.001 (0.7784) | No meaningful change |
| MYOD1 | Muscle differentiation | GSE175495 | 0.003 (0.2905) | 0.006 (0.0010) | Significance changed |
| MYOD1 | Muscle differentiation | GSE226646 | 0.003 (0.2905) | 0.003 (0.3574) | No meaningful change |
| MYOD1 | Muscle differentiation | GSE242202 | 0.003 (0.2905) | 0.004 (0.2054) | No meaningful change |
| MYOD1 | Muscle differentiation | GSE271606 | 0.003 (0.2905) | 0.003 (0.3007) | No meaningful change |
| MSTN | Negative regulator | GSE111010 | 0.002 (0.7230) | -0.002 (0.5267) | No meaningful change |
| MSTN | Negative regulator | GSE111016 | 0.002 (0.7230) | 0.000 (0.9330) | No meaningful change |
| MSTN | Negative regulator | GSE125892 | 0.002 (0.7230) | 0.001 (0.8876) | No meaningful change |
| MSTN | Negative regulator | GSE167186 | 0.002 (0.7230) | 0.005 (0.3895) | No meaningful change |
| MSTN | Negative regulator | GSE175495 | 0.002 (0.7230) | 0.004 (0.4356) | No meaningful change |
| MSTN | Negative regulator | GSE200398 | 0.002 (0.7230) | 0.004 (0.4849) | No meaningful change |
| MSTN | Negative regulator | GSE208639 | 0.002 (0.7230) | 0.001 (0.7791) | No meaningful change |
| MSTN | Negative regulator | GSE287342 | 0.002 (0.7230) | 0.001 (0.8009) | No meaningful change |
| ACTA1 | Structural | GSE111010 | -0.003 (0.0475) | -0.004 (0.0041) | No meaningful change |
| ACTA1 | Structural | GSE111016 | -0.003 (0.0475) | -0.003 (0.0187) | No meaningful change |
| ACTA1 | Structural | GSE113165 | -0.003 (0.0475) | -0.003 (0.0238) | No meaningful change |
| ACTA1 | Structural | GSE125892 | -0.003 (0.0475) | -0.003 (0.0436) | No meaningful change |
| ACTA1 | Structural | GSE144304 | -0.003 (0.0475) | -0.003 (0.0510) | Significance changed |
| ACTA1 | Structural | GSE157712 | -0.003 (0.0475) | -0.003 (0.0231) | No meaningful change |
| ACTA1 | Structural | GSE159217 | -0.003 (0.0475) | -0.004 (<0.0001) | No meaningful change |
| ACTA1 | Structural | GSE167186 | -0.003 (0.0475) | 0.004 (0.3597) | Significance and direction of association changed |
| ACTA1 | Structural | GSE175495 | -0.003 (0.0475) | 0.004 (0.3540) | Significance and direction of association changed |
| ACTA1 | Structural | GSE181107 | -0.003 (0.0475) | -0.003 (0.0532) | Significance changed |
| ACTA1 | Structural | GSE200398 | -0.003 (0.0475) | -0.003 (0.0252) | No meaningful change |
| ACTA1 | Structural | GSE208639 | -0.003 (0.0475) | -0.003 (0.0467) | No meaningful change |
| ACTA1 | Structural | GSE226646 | -0.003 (0.0475) | -0.003 (0.0855) | Significance changed |
| ACTA1 | Structural | GSE242202 | -0.003 (0.0475) | -0.003 (0.0750) | Significance changed |
| ACTA1 | Structural | GSE271606 | -0.003 (0.0475) | -0.003 (0.0600) | Significance changed |
| ACTA1 | Structural | GSE287342 | -0.003 (0.0475) | -0.005 (<0.0001) | No meaningful change |
| MYH1 | Structural | GSE111010 | -0.008 (0.1515) | -0.009 (0.1069) | No meaningful change |
| MYH1 | Structural | GSE111016 | -0.008 (0.1515) | -0.009 (0.0901) | No meaningful change |
| MYH1 | Structural | GSE125892 | -0.008 (0.1515) | -0.008 (0.1131) | No meaningful change |
| MYH1 | Structural | GSE167186 | -0.008 (0.1515) | -0.006 (0.6483) | No meaningful change |
| MYH1 | Structural | GSE175495 | -0.008 (0.1515) | -0.008 (0.1621) | No meaningful change |
| MYH1 | Structural | GSE226646 | -0.008 (0.1515) | -0.005 (0.3784) | No meaningful change |
| MYH1 | Structural | GSE242202 | -0.008 (0.1515) | -0.007 (0.1713) | No meaningful change |
| MYH1 | Structural | GSE271606 | -0.008 (0.1515) | -0.007 (0.1491) | No meaningful change |
| MYH2 | Structural | GSE111010 | -0.005 (0.0050) | -0.005 (0.0041) | No meaningful change |
| MYH2 | Structural | GSE111016 | -0.005 (0.0050) | -0.005 (0.0060) | No meaningful change |
| MYH2 | Structural | GSE113165 | -0.005 (0.0050) | -0.005 (0.0076) | No meaningful change |
| MYH2 | Structural | GSE125892 | -0.005 (0.0050) | -0.005 (0.0578) | Significance changed |
| MYH2 | Structural | GSE144304 | -0.005 (0.0050) | -0.005 (0.0369) | No meaningful change |
| MYH2 | Structural | GSE157712 | -0.005 (0.0050) | -0.005 (0.0054) | No meaningful change |
| MYH2 | Structural | GSE159217 | -0.005 (0.0050) | -0.006 (0.0008) | No meaningful change |
| MYH2 | Structural | GSE167186 | -0.005 (0.0050) | -0.004 (0.0523) | Significance changed |
| MYH2 | Structural | GSE175495 | -0.005 (0.0050) | -0.004 (0.0158) | No meaningful change |
| MYH2 | Structural | GSE181107 | -0.005 (0.0050) | -0.005 (0.0060) | No meaningful change |
| MYH2 | Structural | GSE200398 | -0.005 (0.0050) | -0.005 (0.0054) | No meaningful change |
| MYH2 | Structural | GSE208639 | -0.005 (0.0050) | -0.005 (0.0055) | No meaningful change |
| MYH2 | Structural | GSE226646 | -0.005 (0.0050) | -0.005 (0.0063) | No meaningful change |
| MYH2 | Structural | GSE242202 | -0.005 (0.0050) | -0.005 (0.0034) | No meaningful change |
| MYH2 | Structural | GSE271606 | -0.005 (0.0050) | -0.005 (0.0044) | No meaningful change |
| MYH2 | Structural | GSE287342 | -0.005 (0.0050) | -0.006 (0.0011) | No meaningful change |
| MYH7 | Structural | GSE111010 | 0.003 (0.4490) | 0.003 (0.4689) | No meaningful change |
| MYH7 | Structural | GSE111016 | 0.003 (0.4490) | 0.004 (0.1960) | No meaningful change |
| MYH7 | Structural | GSE113165 | 0.003 (0.4490) | 0.003 (0.4518) | No meaningful change |
| MYH7 | Structural | GSE125892 | 0.003 (0.4490) | 0.003 (0.4558) | No meaningful change |
| MYH7 | Structural | GSE144304 | 0.003 (0.4490) | 0.004 (0.3235) | No meaningful change |
| MYH7 | Structural | GSE157712 | 0.003 (0.4490) | 0.003 (0.4223) | No meaningful change |
| MYH7 | Structural | GSE159217 | 0.003 (0.4490) | 0.004 (0.4099) | No meaningful change |
| MYH7 | Structural | GSE167186 | 0.003 (0.4490) | 0.000 (0.8287) | No meaningful change |
| MYH7 | Structural | GSE175495 | 0.003 (0.4490) | 0.002 (0.6693) | No meaningful change |
| MYH7 | Structural | GSE181107 | 0.003 (0.4490) | 0.003 (0.3737) | No meaningful change |
| MYH7 | Structural | GSE200398 | 0.003 (0.4490) | 0.003 (0.4566) | No meaningful change |
| MYH7 | Structural | GSE208639 | 0.003 (0.4490) | 0.003 (0.4036) | No meaningful change |
| MYH7 | Structural | GSE226646 | 0.003 (0.4490) | 0.001 (0.7637) | No meaningful change |
| MYH7 | Structural | GSE242202 | 0.003 (0.4490) | 0.002 (0.5626) | No meaningful change |
| MYH7 | Structural | GSE271606 | 0.003 (0.4490) | 0.003 (0.4893) | No meaningful change |
| MYH7 | Structural | GSE287342 | 0.003 (0.4490) | 0.002 (0.5850) | No meaningful change |
| PPARGC1A | Mitochondrial regulator | GSE111010 | -0.005 (<0.0001) | -0.006 (<0.0001) | No meaningful change |
| PPARGC1A | Mitochondrial regulator | GSE111016 | -0.005 (<0.0001) | -0.005 (<0.0001) | No meaningful change |
| PPARGC1A | Mitochondrial regulator | GSE113165 | -0.005 (<0.0001) | -0.005 (<0.0001) | No meaningful change |
| PPARGC1A | Mitochondrial regulator | GSE125892 | -0.005 (<0.0001) | -0.005 (<0.0001) | No meaningful change |
| PPARGC1A | Mitochondrial regulator | GSE144304 | -0.005 (<0.0001) | -0.006 (<0.0001) | No meaningful change |
| PPARGC1A | Mitochondrial regulator | GSE157712 | -0.005 (<0.0001) | -0.005 (<0.0001) | No meaningful change |
| PPARGC1A | Mitochondrial regulator | GSE159217 | -0.005 (<0.0001) | -0.005 (<0.0001) | No meaningful change |
| PPARGC1A | Mitochondrial regulator | GSE167186 | -0.005 (<0.0001) | -0.005 (<0.0001) | No meaningful change |
| PPARGC1A | Mitochondrial regulator | GSE175495 | -0.005 (<0.0001) | -0.006 (<0.0001) | No meaningful change |
| PPARGC1A | Mitochondrial regulator | GSE181107 | -0.005 (<0.0001) | -0.005 (<0.0001) | No meaningful change |
| PPARGC1A | Mitochondrial regulator | GSE200398 | -0.005 (<0.0001) | -0.005 (<0.0001) | No meaningful change |
| PPARGC1A | Mitochondrial regulator | GSE208639 | -0.005 (<0.0001) | -0.005 (<0.0001) | No meaningful change |
| PPARGC1A | Mitochondrial regulator | GSE226646 | -0.005 (<0.0001) | -0.006 (<0.0001) | No meaningful change |
| PPARGC1A | Mitochondrial regulator | GSE242202 | -0.005 (<0.0001) | -0.006 (<0.0001) | No meaningful change |
| PPARGC1A | Mitochondrial regulator | GSE271606 | -0.005 (<0.0001) | -0.005 (<0.0001) | No meaningful change |
| PPARGC1A | Mitochondrial regulator | GSE287342 | -0.005 (<0.0001) | -0.006 (<0.0001) | No meaningful change |
| TFAM | Mitochondrial regulator | GSE111010 | 0.001 (0.7105) | 0.001 (0.7421) | No meaningful change |
| TFAM | Mitochondrial regulator | GSE111016 | 0.001 (0.7105) | 0.001 (0.7733) | No meaningful change |
| TFAM | Mitochondrial regulator | GSE113165 | 0.001 (0.7105) | 0.001 (0.7747) | No meaningful change |
| TFAM | Mitochondrial regulator | GSE125892 | 0.001 (0.7105) | 0.002 (0.4596) | No meaningful change |
| TFAM | Mitochondrial regulator | GSE144304 | 0.001 (0.7105) | 0.001 (0.8192) | No meaningful change |
| TFAM | Mitochondrial regulator | GSE157712 | 0.001 (0.7105) | 0.002 (0.6692) | No meaningful change |
| TFAM | Mitochondrial regulator | GSE159217 | 0.001 (0.7105) | 0.001 (0.7679) | No meaningful change |
| TFAM | Mitochondrial regulator | GSE167186 | 0.001 (0.7105) | 0.002 (0.6736) | No meaningful change |
| TFAM | Mitochondrial regulator | GSE175495 | 0.001 (0.7105) | 0.002 (0.6763) | No meaningful change |
| TFAM | Mitochondrial regulator | GSE181107 | 0.001 (0.7105) | 0.000 (0.7513) | No meaningful change |
| TFAM | Mitochondrial regulator | GSE200398 | 0.001 (0.7105) | 0.001 (0.7094) | No meaningful change |
| TFAM | Mitochondrial regulator | GSE208639 | 0.001 (0.7105) | 0.002 (0.6165) | No meaningful change |
| TFAM | Mitochondrial regulator | GSE226646 | 0.001 (0.7105) | 0.001 (0.7812) | No meaningful change |
| TFAM | Mitochondrial regulator | GSE242202 | 0.001 (0.7105) | 0.000 (0.8998) | No meaningful change |
| TFAM | Mitochondrial regulator | GSE271606 | 0.001 (0.7105) | 0.001 (0.6264) | No meaningful change |
| TFAM | Mitochondrial regulator | GSE287342 | 0.001 (0.7105) | 0.002 (0.6198) | No meaningful change |
| TRIM63 | Ubiquitin ligase | GSE111010 | 0.002 (0.1015) | 0.002 (0.0976) | No meaningful change |
| TRIM63 | Ubiquitin ligase | GSE111016 | 0.002 (0.1015) | 0.002 (0.0923) | No meaningful change |
| TRIM63 | Ubiquitin ligase | GSE113165 | 0.002 (0.1015) | 0.002 (0.2493) | No meaningful change |
| TRIM63 | Ubiquitin ligase | GSE125892 | 0.002 (0.1015) | 0.002 (0.1167) | No meaningful change |
| TRIM63 | Ubiquitin ligase | GSE144304 | 0.002 (0.1015) | 0.002 (0.3809) | No meaningful change |
| TRIM63 | Ubiquitin ligase | GSE157712 | 0.002 (0.1015) | 0.002 (0.1039) | No meaningful change |
| TRIM63 | Ubiquitin ligase | GSE159217 | 0.002 (0.1015) | 0.002 (0.3013) | No meaningful change |
| TRIM63 | Ubiquitin ligase | GSE167186 | 0.002 (0.1015) | 0.003 (0.0071) | Significance changed |
| TRIM63 | Ubiquitin ligase | GSE175495 | 0.002 (0.1015) | 0.003 (0.0163) | Significance changed |
| TRIM63 | Ubiquitin ligase | GSE181107 | 0.002 (0.1015) | 0.002 (0.1259) | No meaningful change |
| TRIM63 | Ubiquitin ligase | GSE200398 | 0.002 (0.1015) | 0.002 (0.1161) | No meaningful change |
| TRIM63 | Ubiquitin ligase | GSE208639 | 0.002 (0.1015) | 0.002 (0.0987) | No meaningful change |
| TRIM63 | Ubiquitin ligase | GSE226646 | 0.002 (0.1015) | 0.001 (0.2500) | No meaningful change |
| TRIM63 | Ubiquitin ligase | GSE242202 | 0.002 (0.1015) | 0.002 (0.0782) | No meaningful change |
| TRIM63 | Ubiquitin ligase | GSE271606 | 0.002 (0.1015) | 0.002 (0.1243) | No meaningful change |
| TRIM63 | Ubiquitin ligase | GSE287342 | 0.002 (0.1015) | 0.003 (0.0313) | Significance changed |
| LC3B | Autophagy | GSE111010 | -0.003 (0.5719) | -0.006 (0.2903) | No meaningful change |
| LC3B | Autophagy | GSE111016 | -0.003 (0.5719) | -0.004 (0.4806) | No meaningful change |
| LC3B | Autophagy | GSE208639 | -0.003 (0.5719) | -0.001 (0.8250) | No meaningful change |
| LC3B | Autophagy | GSE242202 | -0.003 (0.5719) | -0.003 (0.5789) | No meaningful change |
| LC3B | Autophagy | GSE271606 | -0.003 (0.5719) | -0.002 (0.7251) | No meaningful change |
| LC3B | Autophagy | GSE287342 | -0.003 (0.5719) | -0.001 (0.8153) | No meaningful change |
| STAR | Steroidogenic enzyme | GSE111010 | -0.004 (0.0070) | -0.003 (0.0219) | No meaningful change |
| STAR | Steroidogenic enzyme | GSE111016 | -0.004 (0.0070) | -0.004 (0.0087) | No meaningful change |
| STAR | Steroidogenic enzyme | GSE113165 | -0.004 (0.0070) | -0.004 (0.0223) | No meaningful change |
| STAR | Steroidogenic enzyme | GSE144304 | -0.004 (0.0070) | -0.004 (0.0134) | No meaningful change |
| STAR | Steroidogenic enzyme | GSE157712 | -0.004 (0.0070) | -0.004 (0.0151) | No meaningful change |
| STAR | Steroidogenic enzyme | GSE159217 | -0.004 (0.0070) | -0.003 (0.0828) | Significance changed |
| STAR | Steroidogenic enzyme | GSE175495 | -0.004 (0.0070) | -0.005 (<0.0001) | No meaningful change |
| STAR | Steroidogenic enzyme | GSE181107 | -0.004 (0.0070) | -0.004 (0.0086) | No meaningful change |
| STAR | Steroidogenic enzyme | GSE200398 | -0.004 (0.0070) | -0.004 (0.0106) | No meaningful change |
| STAR | Steroidogenic enzyme | GSE208639 | -0.004 (0.0070) | -0.004 (0.0095) | No meaningful change |
| STAR | Steroidogenic enzyme | GSE242202 | -0.004 (0.0070) | -0.004 (0.0079) | No meaningful change |
| STAR | Steroidogenic enzyme | GSE271606 | -0.004 (0.0070) | -0.004 (0.0061) | No meaningful change |
| STAR | Steroidogenic enzyme | GSE287342 | -0.004 (0.0070) | -0.004 (0.0117) | No meaningful change |
| TSPO | Steroidogenic enzyme | GSE111010 | 0.000 (0.8558) | 0.000 (0.8820) | No meaningful change |
| TSPO | Steroidogenic enzyme | GSE111016 | 0.000 (0.8558) | 0.000 (0.8434) | No meaningful change |
| TSPO | Steroidogenic enzyme | GSE113165 | 0.000 (0.8558) | 0.000 (0.9630) | No meaningful change |
| TSPO | Steroidogenic enzyme | GSE125892 | 0.000 (0.8558) | 0.000 (0.9755) | No meaningful change |
| TSPO | Steroidogenic enzyme | GSE144304 | 0.000 (0.8558) | -0.001 (0.6286) | No meaningful change |
| TSPO | Steroidogenic enzyme | GSE157712 | 0.000 (0.8558) | 0.000 (0.7202) | No meaningful change |
| TSPO | Steroidogenic enzyme | GSE159217 | 0.000 (0.8558) | -0.001 (0.4645) | No meaningful change |
| TSPO | Steroidogenic enzyme | GSE167186 | 0.000 (0.8558) | 0.001 (0.3572) | No meaningful change |
| TSPO | Steroidogenic enzyme | GSE175495 | 0.000 (0.8558) | 0.000 (0.9888) | No meaningful change |
| TSPO | Steroidogenic enzyme | GSE181107 | 0.000 (0.8558) | 0.000 (0.7898) | No meaningful change |
| TSPO | Steroidogenic enzyme | GSE200398 | 0.000 (0.8558) | 0.000 (0.8687) | No meaningful change |
| TSPO | Steroidogenic enzyme | GSE208639 | 0.000 (0.8558) | 0.000 (0.8815) | No meaningful change |
| TSPO | Steroidogenic enzyme | GSE226646 | 0.000 (0.8558) | 0.000 (0.9465) | No meaningful change |
| TSPO | Steroidogenic enzyme | GSE242202 | 0.000 (0.8558) | 0.000 (0.7684) | No meaningful change |
| TSPO | Steroidogenic enzyme | GSE271606 | 0.000 (0.8558) | 0.000 (0.7816) | No meaningful change |
| TSPO | Steroidogenic enzyme | GSE287342 | 0.000 (0.8558) | 0.000 (0.8183) | No meaningful change |
| HSD3B2 | Steroidogenic enzyme | GSE111010 | -0.003 (0.0005) | -0.002 (0.0027) | No meaningful change |
| HSD3B2 | Steroidogenic enzyme | GSE111016 | -0.003 (0.0005) | -0.002 (0.0026) | No meaningful change |
| HSD3B2 | Steroidogenic enzyme | GSE113165 | -0.003 (0.0005) | -0.003 (<0.0001) | No meaningful change |
| HSD3B2 | Steroidogenic enzyme | GSE125892 | -0.003 (0.0005) | -0.003 (0.0005) | No meaningful change |
| HSD3B2 | Steroidogenic enzyme | GSE144304 | -0.003 (0.0005) | -0.003 (0.0295) | No meaningful change |
| HSD3B2 | Steroidogenic enzyme | GSE157712 | -0.003 (0.0005) | -0.002 (0.0009) | No meaningful change |
| HSD3B2 | Steroidogenic enzyme | GSE159217 | -0.003 (0.0005) | -0.002 (0.0442) | No meaningful change |
| HSD3B2 | Steroidogenic enzyme | GSE181107 | -0.003 (0.0005) | -0.003 (0.0003) | No meaningful change |
| HSD3B2 | Steroidogenic enzyme | GSE200398 | -0.003 (0.0005) | -0.003 (0.0002) | No meaningful change |
| HSD3B2 | Steroidogenic enzyme | GSE208639 | -0.003 (0.0005) | -0.003 (0.0004) | No meaningful change |
| HSD3B2 | Steroidogenic enzyme | GSE242202 | -0.003 (0.0005) | -0.003 (0.0004) | No meaningful change |
| HSD3B2 | Steroidogenic enzyme | GSE271606 | -0.003 (0.0005) | -0.003 (0.0004) | No meaningful change |
| HSD3B2 | Steroidogenic enzyme | GSE287342 | -0.003 (0.0005) | -0.002 (0.0021) | No meaningful change |
| CYP17A1 | Steroidogenic enzyme | GSE111010 | -0.001 (0.4402) | -0.001 (0.3066) | No meaningful change |
| CYP17A1 | Steroidogenic enzyme | GSE111016 | -0.001 (0.4402) | 0.000 (0.6228) | No meaningful change |
| CYP17A1 | Steroidogenic enzyme | GSE113165 | -0.001 (0.4402) | -0.001 (0.5953) | No meaningful change |
| CYP17A1 | Steroidogenic enzyme | GSE144304 | -0.001 (0.4402) | -0.001 (0.2961) | No meaningful change |
| CYP17A1 | Steroidogenic enzyme | GSE157712 | -0.001 (0.4402) | -0.001 (0.3640) | No meaningful change |
| CYP17A1 | Steroidogenic enzyme | GSE159217 | -0.001 (0.4402) | -0.001 (0.2730) | No meaningful change |
| CYP17A1 | Steroidogenic enzyme | GSE175495 | -0.001 (0.4402) | -0.001 (0.3680) | No meaningful change |
| CYP17A1 | Steroidogenic enzyme | GSE181107 | -0.001 (0.4402) | 0.000 (0.6077) | No meaningful change |
| CYP17A1 | Steroidogenic enzyme | GSE200398 | -0.001 (0.4402) | -0.001 (0.4285) | No meaningful change |
| CYP17A1 | Steroidogenic enzyme | GSE208639 | -0.001 (0.4402) | -0.001 (0.4113) | No meaningful change |
| CYP17A1 | Steroidogenic enzyme | GSE242202 | -0.001 (0.4402) | -0.001 (0.3408) | No meaningful change |
| CYP17A1 | Steroidogenic enzyme | GSE271606 | -0.001 (0.4402) | 0.000 (0.5889) | No meaningful change |
| CYP17A1 | Steroidogenic enzyme | GSE287342 | -0.001 (0.4402) | 0.000 (0.5731) | No meaningful change |
| SRD5A1 | Steroidogenic enzyme | GSE111010 | -0.001 (0.3043) | -0.001 (0.2388) | No meaningful change |
| SRD5A1 | Steroidogenic enzyme | GSE111016 | -0.001 (0.3043) | -0.002 (0.2044) | No meaningful change |
| SRD5A1 | Steroidogenic enzyme | GSE113165 | -0.001 (0.3043) | -0.002 (0.1941) | No meaningful change |
| SRD5A1 | Steroidogenic enzyme | GSE144304 | -0.001 (0.3043) | 0.000 (0.8700) | No meaningful change |
| SRD5A1 | Steroidogenic enzyme | GSE157712 | -0.001 (0.3043) | -0.001 (0.3052) | No meaningful change |
| SRD5A1 | Steroidogenic enzyme | GSE159217 | -0.001 (0.3043) | -0.001 (0.3001) | No meaningful change |
| SRD5A1 | Steroidogenic enzyme | GSE175495 | -0.001 (0.3043) | 0.000 (0.9361) | No meaningful change |
| SRD5A1 | Steroidogenic enzyme | GSE181107 | -0.001 (0.3043) | -0.001 (0.2962) | No meaningful change |
| SRD5A1 | Steroidogenic enzyme | GSE200398 | -0.001 (0.3043) | -0.001 (0.3387) | No meaningful change |
| SRD5A1 | Steroidogenic enzyme | GSE208639 | -0.001 (0.3043) | -0.001 (0.3720) | No meaningful change |
| SRD5A1 | Steroidogenic enzyme | GSE242202 | -0.001 (0.3043) | -0.001 (0.3442) | No meaningful change |
| SRD5A1 | Steroidogenic enzyme | GSE271606 | -0.001 (0.3043) | -0.001 (0.3150) | No meaningful change |
| SRD5A1 | Steroidogenic enzyme | GSE287342 | -0.001 (0.3043) | -0.001 (0.3290) | No meaningful change |

Supplementary Table 4. Leave-one-out analysis: Summary of influential datasets. *LOO: Leave-one-out*

| Gene | Category | Excluded Dataset | Full Model Beta (p-value) | LOO Model Beta (p-value) | Impact of dataset exclusion |
| --- | --- | --- | --- | --- | --- |
| FKBP4 | Chaperone | GSE111010 | 0.003 (0.0177) | 0.002 (0.0543) | Significance changed |
| FKBP4 | Chaperone | GSE111016 | 0.003 (0.0177) | 0.002 (0.0562) | Significance changed |
| FKBP4 | Chaperone | GSE125892 | 0.003 (0.0177) | 0.003 (0.0527) | Significance changed |
| FKBP4 | Chaperone | GSE144304 | 0.003 (0.0177) | 0.002 (0.1119) | Significance changed |
| FKBP4 | Chaperone | GSE159217 | 0.003 (0.0177) | 0.002 (0.0752) | Significance changed |
| FKBP4 | Chaperone | GSE287342 | 0.003 (0.0177) | 0.002 (0.0514) | Significance changed |
| FKBP5 | Chaperone | GSE144304 | 0.006 (0.0126) | 0.005 (0.0656) | Significance changed |
| FKBP5 | Chaperone | GSE167186 | 0.006 (0.0126) | 0.005 (0.0627) | Significance changed |
| KPNA1 | Nuclear transport | GSE167186 | 0.011 (0.0706) | 0.015 (0.0352) | Significance changed |
| KPNA1 | Nuclear transport | GSE175495 | 0.011 (0.0706) | 0.015 (0.0224) | Significance changed |
| KPNA5 | Nuclear transport | GSE144304 | 0.002 (0.1080) | 0.002 (0.0098) | Significance changed |
| KPNA5 | Nuclear transport | GSE167186 | 0.001 (0.4685) | 0.002 (0.0401) | Significance changed |
| NCOA2 | Coactivator | GSE175495 | -0.002 (0.0120) | -0.002 (0.0756) | Significance changed |
| MED1 | Mediator complex | GSE175495 | -0.002 (0.0120) | -0.002 (0.1668) | Significance changed |
| SMARCA4 | Chromatin remodeler | GSE111016 | -0.002 (0.0120) | -0.002 (0.0832) | Significance changed |
| SMARCA4 | Chromatin remodeler | GSE144304 | -0.002 (0.0120) | -0.001 (0.1425) | Significance changed |
| SMARCA4 | Chromatin remodeler | GSE157712 | -0.002 (0.0120) | -0.002 (0.1095) | Significance changed |
| SMARCA4 | Chromatin remodeler | GSE242202 | -0.002 (0.0120) | -0.001 (0.1186) | Significance changed |
| SMARCA4 | Chromatin remodeler | GSE287342 | -0.002 (0.0120) | -0.002 (0.0781) | Significance changed |
| KDM1A | Chromatin modifier | GSE144304 | -0.002 (0.0120) | -0.002 (0.0839) | Significance changed |
| STUB1 | Ubiquitination | GSE159217 | -0.002 (0.0120) | -0.001 (0.0854) | Significance changed |
| STUB1 | Ubiquitination | GSE175495 | 0.000 (0.9677) | -0.001 (0.0252) | Significance changed |
| UBE2I | SUMOylation | GSE157712 | -0.002 (0.0195) | -0.002 (0.0716) | Significance changed |
| MDM2 | Ubiquitination | GSE144304 | -0.002 (0.0195) | -0.003 (0.0505) | Significance changed |
| MDM2 | Ubiquitination | GSE159217 | -0.001 (0.1833) | -0.002 (0.0459) | Significance changed |
| MDM2 | Ubiquitination | GSE167186 | 0.002 (0.1354) | 0.003 (0.0194) | Significance changed |
| AKT1 | Kinase | GSE113165 | -0.001 (0.1424) | -0.001 (0.0449) | Significance changed |
| AKT1 | Kinase | GSE208639 | 0.003 (0.2891) | 0.006 (0.0010) | Significance changed |
| MAPK1 | Kinase | GSE159217 | -0.001 (0.5949) | -0.003 (0.0473) | Significance changed |
| MYOD1 | Muscle differentiation | GSE175495 | -0.004 (0.0361) | -0.004 (0.1418) | Significance changed |
| ACTA1 | Structural | GSE144304 | -0.004 (0.0361) | -0.005 (0.0784) | Significance changed |
| ACTA1 | Structural | GSE167186 | -0.004 (0.0361) | -0.005 (0.0760) | Significance changed |
| ACTA1 | Structural | GSE175495 | -0.004 (0.0361) | -0.004 (0.1675) | Significance changed |
| ACTA1 | Structural | GSE181107 | 0.002 (0.0440) | 0.002 (0.0728) | Significance changed |
| ACTA1 | Structural | GSE226646 | 0.002 (0.0440) | 0.002 (0.1184) | Significance changed |
| ACTA1 | Structural | GSE242202 | 0.002 (0.0440) | 0.002 (0.0513) | Significance changed |
| ACTA1 | Structural | GSE271606 | 0.002 (0.0440) | 0.002 (0.2096) | Significance changed |
| MYH2 | Structural | GSE125892 | 0.002 (0.0440) | 0.002 (0.0647) | Significance changed |
| MYH2 | Structural | GSE167186 | 0.002 (0.0440) | 0.003 (0.1661) | Significance changed |
| TRIM63 | Ubiquitin ligase | GSE167186 | 0.002 (0.0440) | 0.002 (0.0505) | Significance changed |
| TRIM63 | Ubiquitin ligase | GSE175495 | 0.002 (0.0440) | 0.002 (0.0505) | Significance changed |
| TRIM63 | Ubiquitin ligase | GSE287342 | 0.002 (0.0440) | 0.003 (0.1201) | Significance changed |
| STAR | Steroidogenic enzyme | GSE159217 | 0.002 (0.0440) | 0.002 (0.1194) | Significance changed |
